# Receptor-tethered orthogonal IL-2 enhances regulatory T cell therapy

**DOI:** 10.1101/2025.07.19.665688

**Authors:** Alyssa Indart, Huiyun Lyu, Vinh Q. Nguyen, Wendy Rosenthal, Sophie S. Jang, Yue Chen, Kevin Jude, Leon Su, K. Christopher Garcia, Jeffrey A. Bluestone, Qizhi Tang

## Abstract

Regulatory T cell (Treg) therapy is an emerging platform for controlling immune overactivation. Efficacy of Treg therapy is limited by the poor persistence of infused Tregs due to insufficient IL-2, which is essential for Treg survival and function. IL-2 activates many immune cells besides Tregs, imposing a challenge for the selective provision of IL-2 to infused Tregs. In this study, we found infusions of orthogonal (ortho) IL-2 failed to enhance Tregs engineered with a corresponding orthoIL-2 receptor (IL-2R) in a mouse model of autoimmune diabetes. We then developed a receptor-tethered orthoIL-2 by optimizing the combination of IL-2, IL-2R, and the linker connecting them to achieve autocrine signaling selectively in engineered Tregs. Tregs expressing the tethered orthoIL-2 showed autocrine IL-2 signaling in vitro, enhanced CD25, CTLA-4, and Foxp3 expression, persisted without exogenous IL-2 in vivo, and prevented autoimmune diabetes using as few as 2,000 Tregs. Knocking the tethered orthoIL-2 construct into the Foxp3 locus enabled Treg-specific expression, with the additional benefit of positively reinforcing tethered orthoIL-2 expression through the activation of the Foxp3 gene by enhanced IL-2 signaling. Together, these results illustrate a safe and effective cell-engineering solution for overcoming Tregs’ dependency on exogenous IL-2, thereby achieving superior therapeutic efficacy.

## Introduction

The essential function of regulatory T cells (Tregs) in maintaining immune homeostasis is illustrated by the life-threatening autoimmune syndrome in human and mice with mutations in the *FOXP3* gene, a Treg-lineage-determining transcription factor(Dikiy and Rudensky, 2023; Hori, 2021; Ramsdell and Rudensky, 2020; Ramsdell and Ziegler, 2014; Sakaguchi et al., 2020). Moreover, the development of many more common autoimmune diseases can be attributed to an imbalance between Tregs and autoreactive T cells(Sumida et al., 2024). Treg therapy effectively suppresses a wide range of autoimmune and inflammatory diseases in preclinical models, prompting the ongoing research and development of Treg cell therapies to treat autoimmune, inflammatory, and degenerative diseases in humans(Bluestone et al., 2023; Esensten et al., 2018; Ho et al., 2024; Pilat and Sprent, 2020; Sakaguchi et al., 2020).

Tregs constitutively express CD25, the interleukin 2 (IL-2) receptor α chain (IL-2Rα). This enables Tregs to respond to low concentrations of IL-2 by stabilizing IL-2 interaction with the signal-transducing IL-2Rβ and IL-2Rγ chains of the IL-2R complex(Malek and Castro, 2010). IL-2, a T-cell growth and survival factor produced by activated T cells(Chinen et al., 2016; Malek and Castro, 2010), plays an indispensable role in the development, proliferation, and survival of Tregs(Fontenot et al., 2005; Rubtsov et al., 2010; Setoguchi et al., 2005). IL-2 signaling reinforces Treg lineage identity by increasing Foxp3 expression and sustains high expression of CD25 and CTLA-4 that are important for Treg function(Fontenot et al., 2005; Jamison et al., 2024; Pandiyan et al., 2007). Paradoxically, Tregs do not produce IL-2 and are dependent on exogenous IL-2 produced by conventional T cells (Tconvs). Thus, the IL-2 and IL- 2R axis plays a pivotal role in regulating the dynamics between immunosuppressive Tregs and immune-activating Tconvs(Busse et al., 2010; Feinerman et al., 2010; Fontenot et al., 2005; Jamison et al., 2024; Wong et al., 2021). Genome-wide association studies have revealed linkage of polymorphisms in *IL2* and *IL2RA* gene loci to various autoimmune diseases, including type 1 diabetes, multiple sclerosis, celiac disease, autoimmune hepatitis, and Crohn’s disease(Hartmann et al., 2014; Todd et al., 2007; van Heel et al., 2007), underscoring the importance of IL-2 signaling in immune self-tolerance.

In a phase 1 Treg therapy trial in type 1 diabetes, we observed a loss of 75% the infused Tregs within 3 months of infusion(Bluestone et al., 2015). In a follow-up trial, we found co-infusion of low-dose IL-2 supported the persistence of the infused Tregs (Fig. S1) but also activated cytotoxic T cells(Dong et al., 2021). The narrow therapeutic window between targeting Tregs and off-Treg activities has been an unsolved challenge for IL-2-based approaches for promoting Tregs(Khoryati et al., 2020; Lykhopiy et al., 2023). Thus, insufficient IL-2 limits the persistence of infused Tregs, but low-dose IL-2 is not reliably selective for Tregs to be a viable solution.

An orthogonal (ortho) IL-2 may achieve selective signaling in Tregs engineered with a corresponding orthoIL-2R(Sockolosky et al., 2018). In this study, we used the non-obese diabetes (NOD) mouse model of human type 1 diabetes to test this approach. We have previously shown the efficacy and dose-dependent toxicity of IL-2 therapy in this model(Grinberg-Bleyer et al., 2010; Tang et al., 2008). Moreover, diabetes progression in the NOD mice can be controlled by Tregs(Tang et al., 2004; Tarbell et al., 2004; Yang et al., 2022). We thus used this model to determine if an orthoIL-2-IL-2R system could synergize with therapeutic Tregs. We found that soluble orthoIL-2 was not sufficiently potent or selective to support therapeutic Tregs *in vivo*, which prompted us to develop a cell-autonomous tethered orthoIL-2-IL-2R construct that sustained Treg survival and improved the potency of the Tregs in preventing autoimmune diabetes. Lastly, we show that targeting this orthoIL-2 autocrine construct to the Foxp3 locus improved the safety of the strategy by avoiding off-Treg expression and reinforced Treg lineage stability via a positive feedback loop.

## Results

### Orthogonal IL-2 3A10 was highly selective for orthoIL-2R^+^ Tregs but lacked potency *in vivo*

A mouse orthoIL-2 system has been developed by screening for ligands that preferentially bind to a mutated IL-2Rβ. Among a panel of orthoIL-2s generated, 3A10, is a highly selective for orthoIL-2R with no measurable activity in cells expressing the wtIL-2R(Sockolosky et al., 2018). We thus expressed the orthoIL-2Rβ in NOD Tregs via retroviral transduction and tested effect of 3A10 *in vitro* and *in vivo* (Fig. 1a). NOD Tregs transduced with the orthoIL-2R, but not the empty vector (EV), responded to 3A10 by phosphorylating STAT5 (pSTAT5) with an EC50 of 8,620 IU/ml (Fig. 1b, Fig. S2). In contrast, wtIL-2 induced maximal pSTAT5 in both EV- and orthoIL-2R-transduced Tregs at the lowest concentration tested (100 IU/ml, Fig. 1b). Furthermore, 100,000 IU/ml 3A10 induced proliferation of orthoIL-2R^+^, but not EV-transduced, Tregs (Fig. 1c). Additionally, 3A10 mildly enhanced the function of orthoIL-2R^+^ Tregs in an *in vitro* suppression assay (Fig. 1d).

**Fig. 1.**
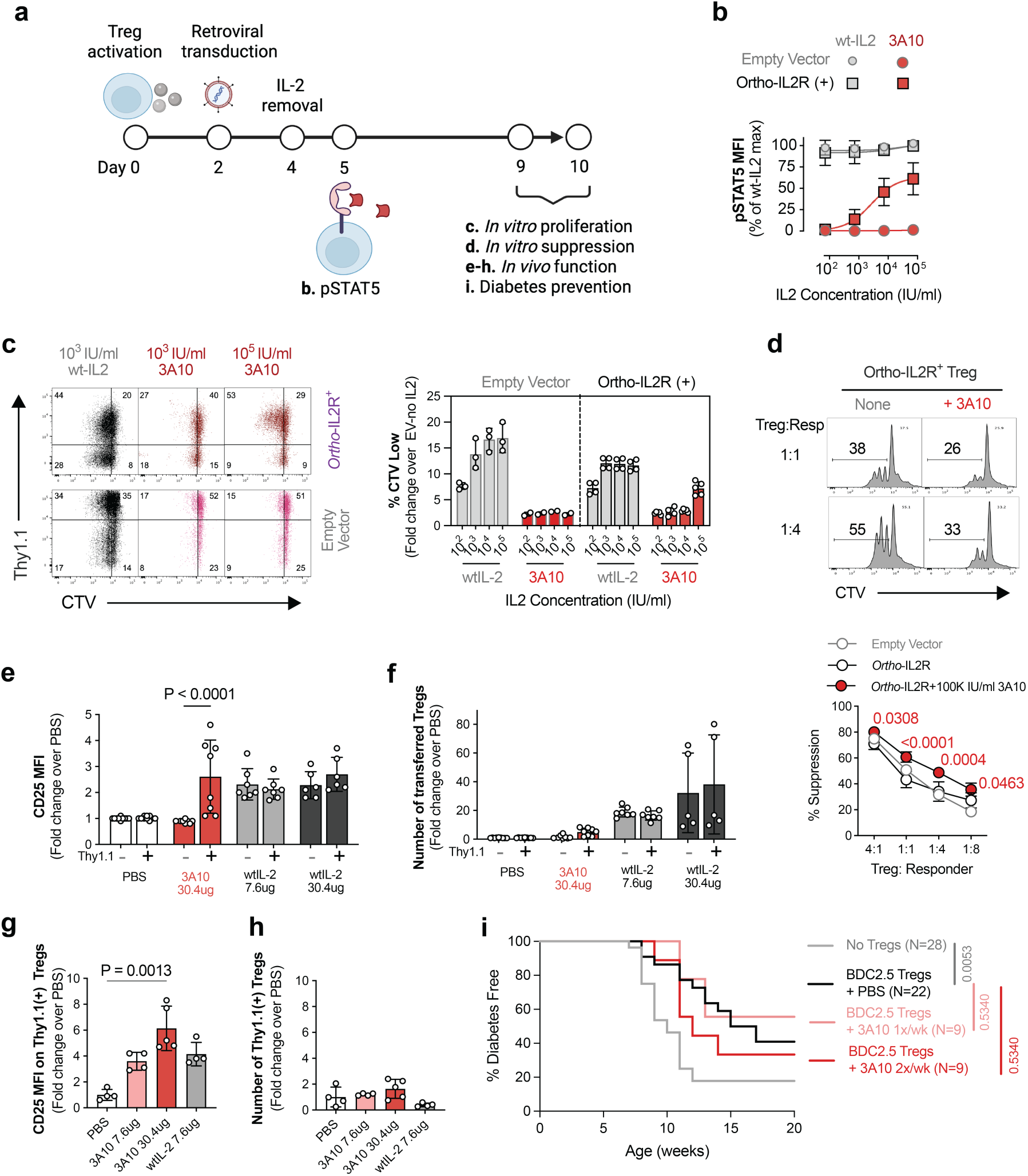
OrthoIL-2 3A10 is a selective but weak agonist for orthoIL-2R. **a**. Experimental workflow for assessing function of 3A10 *in vitro* and *in vivo*. **b.** Dose response of IL-2-induced pSTAT5 in Tregs transduced with empty vector or orthoIL-2Rβ with titrated concentrations of wtIL-2 or orthoIL-2 3A10. **c.** Treg proliferation measured using CTV dilution. **d.** *In vitro* suppression. Ordinary two-way ANOVA followed by Tukey’s multiple comparison post-test was used to determine the statistical significance of the difference observed. P values on the graph are for comparisons between orthoIL-2R^+^ Tregs with or without added 100,000 IU/ml 3A10 at each Treg:Responder ratio. Results in panels **b** to **d** are summaries of 3 independent experiments (means +/- SD, n=3). **e, f.** NSG mice were injected with orthoIL-2R^+^ Tregs and treated daily for 1 week as indicated. CD25 expression (**e**) and number of transferred Tregs **(f)** in the spleens normalized to the means of PBS control are summarized (means +/- SD, n=8-10 mice per group from 3 experiments). Two-way ANOVA followed by Sidak multiple comparison post-test was used to assess statistical significance of the difference between OrthoIL-2R^-^ and OrthoIL-2R^+^ cells under each treatment condition. Only statistically significant P values are listed. **g, h.** NOD.CD28KO mice were injected with BDC2.5 orthoIL-2R^+^ Tregs and treated for 1 week daily. CD25 MFI and number of transferred Tregs in the spleens are summarized (means +/- SD, n=4-5 mice per group pooled from 3 experiments). Kruskal-Wallis test was used to determine the statistical significance from the PBS-treated controls. Only statistically significant P values are listed. **i.** NOD.CD28KO mice were treated with either 2,000 BDC2.5 Treg + PBS or BDC2.5 orthoIL-2R^+^ Tregs + MSA- 3A10 30 μg once or twice per week for 15 weeks. Mice with two readings of blood glucose >250 mg/dL were considered diabetic (n=9-28 mice per group pooled from 2 experiments). Statistical significance was determined using Kaplan-Meier survival analysis. P values were calculated using Mantel-Cox test comparing each condition to the BDC2.5 Treg-treated group.

To test the efficacy of 3A10 on orthoIL-2R^+^ Tregs *in vivo*, we transferred orthoIL-2R^+^ Tregs along with non-transduced Tregs into immuno-deficient, therefore IL-2 deficient, NOD-scid IL-2Rgamma^null^ (NSG) mice followed by daily infusions of mouse serum albumin (MSA)- 3A10 fusion protein, MSA-wtIL-2, or PBS as a control (Fig. S3a). The MSA fusion proteins were used to extend the half-life of 3A10 and wtIL-2 from 5 to 50 hours(Sockolosky et al., 2018). High-dose 3A10 (30.4 μg daily) induced weak proliferation (Fig. S3b) and selectively increased CD25, Foxp3, and CTLA-4 expression on Thy1.1^+^ orthoIL-2R^+^ Tregs (Fig. 1e, Fig. S3c to g). In addition, MSA-3A10 mildly increased the number of Thy1.1^+^ orthoIL-2R^+^ but not Thy1.1^-^ Tregs (Fig. 1f). These data showed that orthoIL-2 3A10 was highly selective for orthoIL-2R^+^ Tregs but only weakly stimulated orthoIL-2R^+^ Tregs *in vitro* and *in vivo*.

To determine the impact of MSA-3A10 on the function of orthoIL-2R^+^ Tregs in lymphoreplete recipients, we investigated the homeostasis and function of orthoIL-2R^+^ in NOD.CD28KO mice. These mice develop diabetes at an accelerated tempo and higher penetrance due to defective Treg development and peripheral homeostasis, thus serving as a robust model for evaluating Treg function in autoimmune diabetes *in vivo*(Mahne et al., 2015; Obarorakpor et al., 2023; Salomon et al., 2000; Tang et al., 2004). In these experiments, we used islet antigen-specific Tregs from BDC2.5 TCR transgenic mice (referred to as BDC2.5 Tregs hereafter) because of their established efficacy in this model (Fig. S4a). The effects of MSA- 3A10 were muted in NOD.CD28KO mice when compared to those observed in NSG mice.

Despite the significant increase in CD25 expression on transferred orthoIL-2R^+^ Tregs, their total cell numbers were not different from PBS-treated controls in the spleen (Fig. 1g, h). In the pancreatic LN (pancLN) where the cognate antigen for the BDC2.5 TCR was present, no increase in either CD25 expression or the number of orthoIL-2R^+^ Tregs was seen (Fig. S4b, c). The effect of MSA-3A10 was selective for orthoIL-2R^+^ Tregs since no change in CD25 expression was observed in the spleen or pancLN on endogenous Tregs, CD4^+^, and CD8^+^ Tconvs (Fig. S4d-i). These results show that MSA-3A10 had limited ability to promote proliferation of orthoIL-2R^+^ Tregs in NOD.CD28KO mice.

We next determined if MSA-3A10 could enhance the efficacy of Tregs in preventing autoimmune diabetes. Previously, we have shown that 50,000 to 150,000 BDC2.5 Tregs prevented diabetes in 100% of NOD.CD28KO recipients(Spence et al., 2018; Tang et al., 2004). A dose titration experiment showed that 10,000 BDC2.5 Tregs were similarly effective whereas 2,000 BDC2.5 Tregs were partially protective (Fig. S4j). We thus infused mice with 2,000 BDC2.5^+^ orthoIL-2R^+^ Tregs and assessed the impact of MSA-3A10 supplements. No improvement in diabetes-free survival was observed (Fig. 1i). Taken together, these data indicate 3A10 was a highly selective ligand for the orthoIL-2R, but its weak activity limited its *in vivo* efficacy in potentiating orthoIL-2R^+^ Tregs.

### Orthogonal IL-2 1G12 was more potent but lacked selectivity for orthoIL-2R^+^ Tregs

Realizing the limitation of 3A10’s low affinity, we explored the performance of another orthoIL- 2, 1G12, that had higher affinity for orthoIL-2R^+^ but could also signal via wtIL-2R at high concentrations(Sockolosky et al., 2018). 1G12 was more potent than 3A10 with pSTAT5 EC_50_ of 100 IU/ml in orthoIL-2R^+^ Tregs. It also showed agonist activity in EV-transduced wtIL-2R- expressing Tregs with pSTAT5 EC_50_ of 1,714 IU/ml (Fig. 2a, Fig. S5a). 1G12 induced *in vitro* proliferation of orthoIL-2R^+^ Tregs but not in EV-transduced Tregs at low concentrations (1,000 IU/ml) (Fig. 2b). At concentrations above 10,000 IU/ml, 1G12 induced proliferation of both orthoIL-2R^+^ and EV-transduced Tregs (Fig. 2b). Similarly, 1G12’s effect on *in vitro* suppression was dose dependent. At 1,000 IU/ml, no change in the suppressive activity of orthoIL-2R^+^ Tregs was observed; slightly enhanced suppression was seen at 10,000 IU/ml of 1G12, whereas significantly impaired suppression was seen with 100,000 IU/ml of 1G12 at 1:8 Treg:Tconv ratio (Fig. 2c), likely due to its activation of responder T cells in the assay.

**Fig. 2.**
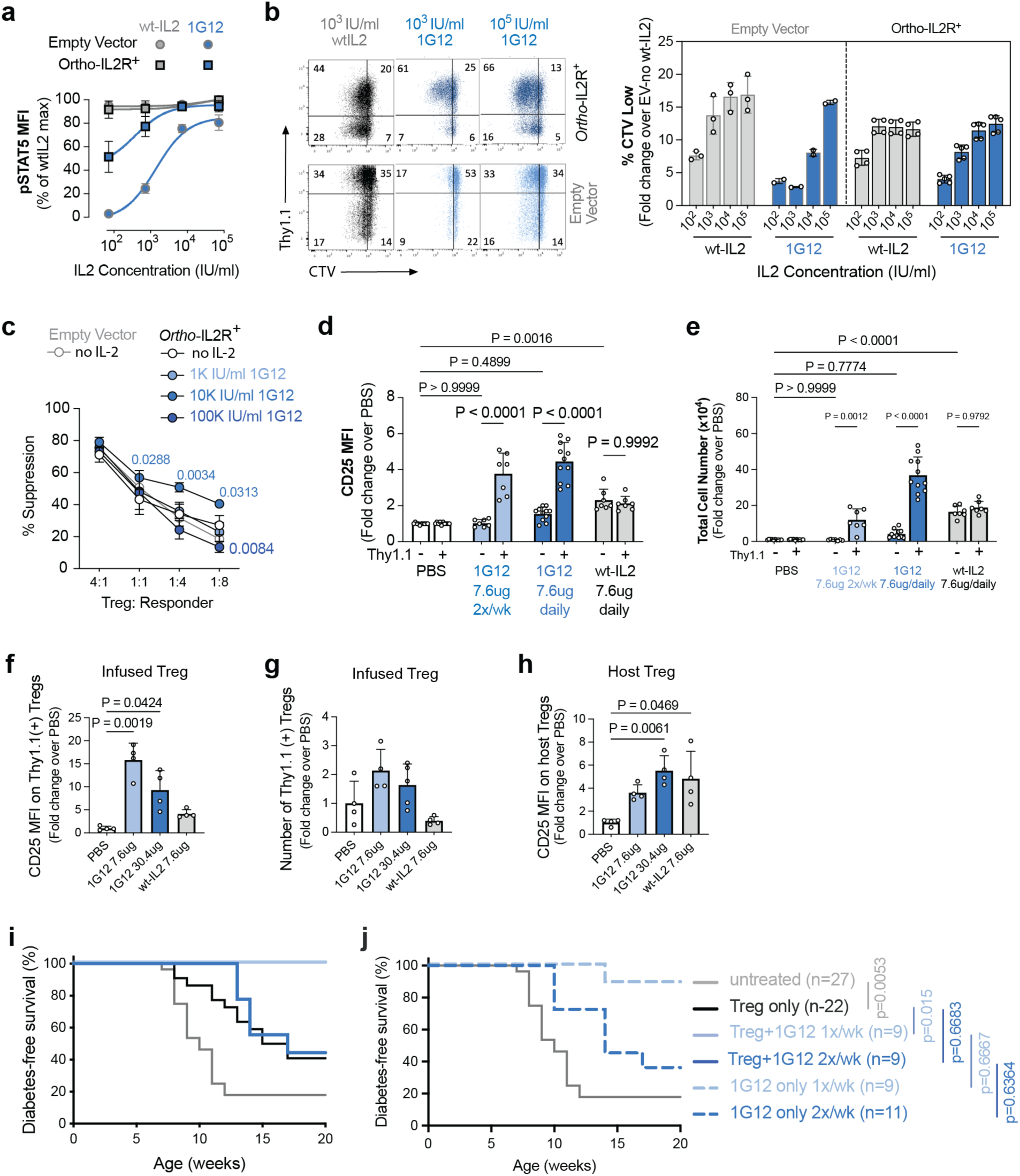
OrthoIL-2 1G12 has higher potency but cross-reacts on wtIL-2R. **a.** Dose response of IL-2-induced pSTAT5 in Tregs transduced with empty vector or orthoIL-2Rβ. **b.** Treg proliferation measured using CTV dilution. (**c.**) *In vitro* suppression assay. Results shown are summaries of 3 experimental replicates (means +/- SD, n=3). Results shown in panels **a-c** are summaries of 3 independent experiments (means +/- SD, n=3). Ordinary two-way ANOVA followed by Tukey’s multiple comparison post-test was used to determine the statistical significance of the difference observed. P values on the graph are for comparisons with orthoIL-2R+ Tregs without addition of 1G12 at each Treg:Responder ratio. Only P values <0.05 are listed. **d, e.** NSG mice were injected with orthoIL-2R^+^ Tregs and treated daily for 1 week as indicated. CD25 expression (**d**) and number of transferred Tregs (**e**) in the spleens normalized to the means of PBS control are summarized (means +/- SD, n=8-10 mice per group from 3 experiments). Two-way ANOVA followed by Tukey’s multiple comparison post-test was used to assess statistical significance between experimental groups. **f.** Experiments for assessing function of 1G12 in NOD.CD28KO mice. **g-i.** NOD.CD28KO mice were injected with BDC2.5 orthoIL-2R^+^ Tregs and treated for 1 week daily as indicated. CD25 expression by transferred Tregs (**g**) number of transferred Tregs (**h**), and CD25 MFI on host Tregs (**i**) in the spleens are summarized (means +/- SD, n=4-5 mice per group from 3 experiments). Kruskal-Wallis test was used to determine the statistical significance from the PBS-treated controls. Only statistically significant P values are shown. **j, k**. NOD.CD28KO mice were treated with PBS or 7.6 μg MSA-1G12 for 15 weeks as indicated with (**j**) or without (**k**) 2,000 orthoIL-2R^+^ BDC2.5 Tregs. Mice with two readings of blood glucose >250 mg/dL were considered diabetic (n=9-28 mice per group pooled from 2 experiments). Statistical significance was determined using Kaplan- Meier survival analysis. P values were calculated using Mantel-Cox test comparing each condition to BDC2.5 Treg- treated group and comparing the same dose of MSA-1G12 with or without BDC2.5 Tregs.

In NSG mice, orthoIL-2R^+^ Tregs proliferated comparably when the mice received 7.6 or 30.4 μg of MSA-1G12 or MSA-wtIL-2 (Fig. S5b, c), suggesting that daily infusion of 7.6 μg of either IL-2 form was saturating. We thus eliminated the 30.4 μg dose in follow-up experiments and added 7.6 μg twice per week dosing. We observed that daily and twice per week MSA-1G12 were similarly effective in inducing CD25, Foxp3, and CTLA-4 expression on orthoIL-2R^+^ Tregs (Fig. 2d and Fig. S5d to f), but daily dosing induced a significantly greater increase in the number of orthoIL-2R^+^ Tregs when compared to twice per week dosing (Fig. 2e). Moreover, neither dose of MSA-1G12 significantly changed CD25 expression or the cell numbers of Thy1.1^-^ orthoIL-2R^-^ cells in these short-term experiments (Fig. 2d and e).

In lymphoreplete NOD.CD28KO recipients (Fig. S6a), 7.6 μg and 30.4 μg daily MSA- 1G12 significantly increased the expression of CD25 on transferred orthoIL-2R^+^ Tregs in the spleen (Fig. 2f), pancLN (Fig. S6b), and pancreatic islets (Fig. S6c). However, the total numbers of infused orthoIL-2R^+^ Tregs in the spleens (Fig. 2g) and pancLNs (Fig. S6b) were not significantly more than PBS-treated controls. Moreover, we observed a significant increase in CD25 expression on host Treg in the spleen (Fig. 2h) and pancLN (Fig. S6b) with 30.4 μg MSA- 1G12, demonstrating cross reactivity on host cells at this high dose. The expression of CD25 on host CD4^+^ or CD8^+^ Tconvs in the spleens and pancLN were largely unchanged (Fig. S6d and e).

Lastly, NOD.CD28KO mice treated with 2,000 BDC2.5 orthoIL-2R^+^ Tregs followed by 7.6 μg of MSA-1G12 once a week were 100% protected from diabetes, but MSA-1G12 twice a week did not differ from Treg alone (Fig. 2i). Since MSA-1G12 could act on endogenous Tregs and Tconvs, we treated NOD.CD28KO mice once or twice a week with 7.6 μg MSA-1G12 without infusion of Tregs. The results mirrored those with Treg infusion (Fig. 3j). Thus, 1G12 controlled diabetes independent of the infused Tregs and its effect was dose dependent, similar to previous observations with wtIL-2(Grinberg-Bleyer et al., 2010; Tang et al., 2008). These data together show the limited utility of 1G12 as an adjunct therapy to therapeutic Tregs.

**Fig. 3.**
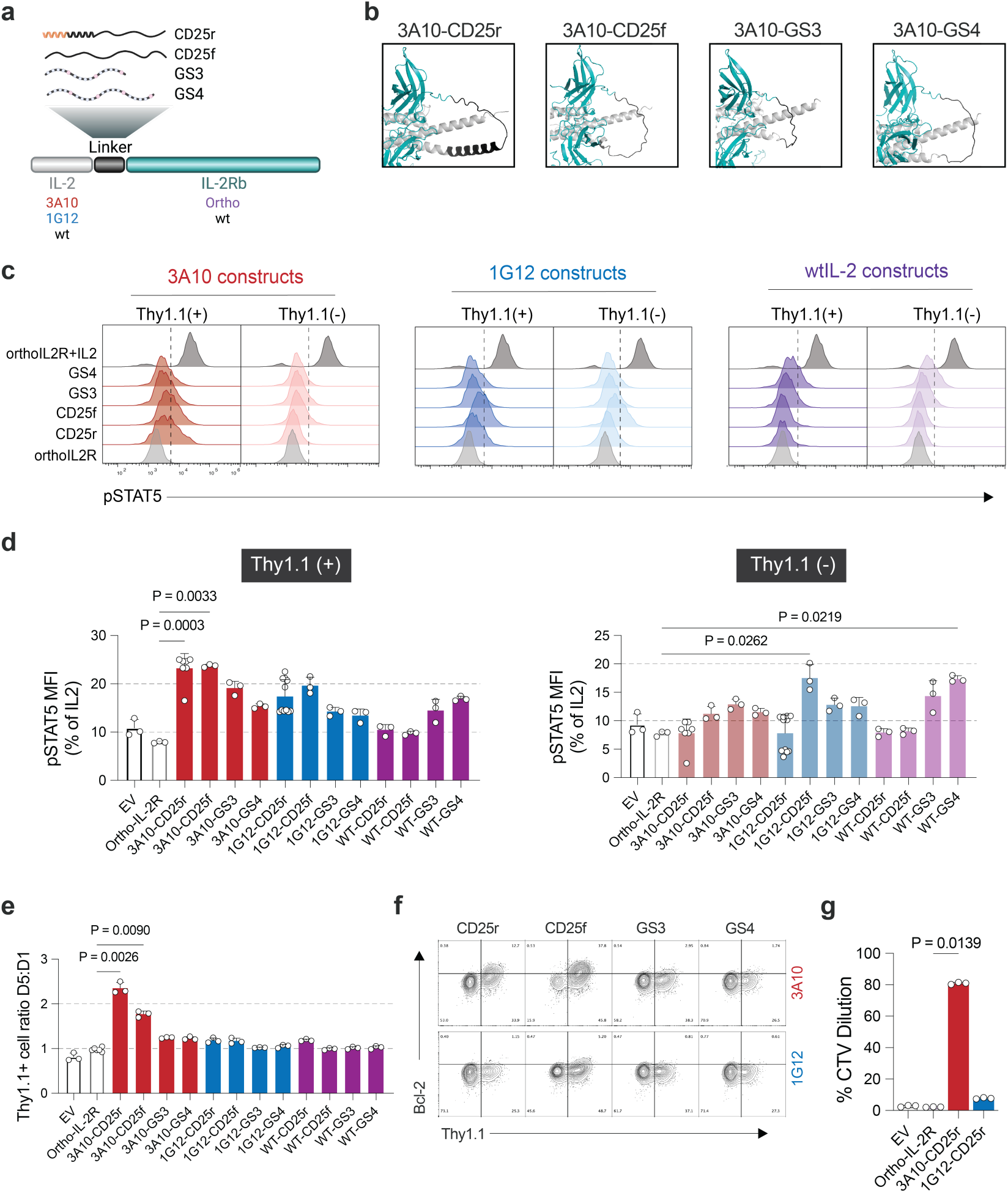
**Engineering a tethered orthoIL-2-IL-2R**β **system. a.** Schematic of the tethered IL-2-IL-2R components tested. **b.** Visualization of the four linkers used to tether orthoIL-2 3A10 to orthoIL-2R using Pymol. **c, d.** Flow cytometric plots of pSTAT5 in transduced Tregs (Thy1.1^+^, left) or not (Thy1.1^-^, right) with various constructs indicated. Representative flow plots are shown. **c.** Results were repeated in 3 independent experiments. **d.** Data were expressed as percentages of pSTAT5 MFI relative to the that induced in orthoIL- 2R-transduced Tregs stimulated with 100 IU/ml IL-2. **e.** Enrichment of Thy1.1^+^ Tregs on day 5 versus day 1 after exogenous IL-2 withdrawal. Results shown are a summary of 3 independent experiments (mean+/- SD, n=3-6). **f.** Representative flow cytometric plots of Bcl-2 expression in Tregs transduced with various tethered 3A10 and 1G12 constructs. The results shown are representative of 3 independent experiments. **g.** On day 5 after activation, Thy1.1^+^ Tregs were purified, labelled with CTV, and cultured without TCR activation or exogenous IL-2. CTV dilution was measured on day 14. For panels d, e, and g, Kruskal-Wallis test was used to determine the statistical significance when compared to the orthoIL-2R controls. Only statistically significant P values are shown.

### Potency and selectivity of tethered orthoIL-2-IL-2Rβ system depended on the ligand and the linker

We hypothesized that physically tethering the orthoIL-2 to the orthoIL-2R would improve the potency and selectivity of the orthoIL-2-IL-2R system. We thus designed 4 linkers containing 15-26 amino acids, which should be sufficient to span the distance of 41 angstroms between the C-terminus of the IL-2 and the N-terminus of the IL-2Rβ (Fig. 3a). The first linker was a 26- amino-acid peptide from the extracellular domain of CD25(Jounaidi et al., 2017). The linker contains a short rigid alpha helix followed by a flexible disordered domain of CD25, which we named CD25-rigid (CD25r). The second linker was a 26-amino-acid peptide entirely from the disordered region of the CD25 extracellular domain to give the linker more flexibility. We termed this linker CD25-flexible (CD25f). The other two linkers were 3 or 4 repeats of GGGGS, named GS3 and GS4, respectively. We used each of these linkers to tether 3A10 or 1G12 to orthoIL-2Rβ. We also generated 4 analogous wtIL-2 tethered to wtIL-2Rβ with these linkers to test if tethering IL-2 would be sufficient to restrict IL-2 signaling to the engineered cells to obviate the need for orthogonality. Alpha Fold models showed that the side chains of tethered 3A10 were 3 to 6 angstroms away from the orthoIL-2Rβ binding pocket (Fig. 3b, Fig. S7a) and there was no steric hindrance from the linkers (Fig. S7b).

Tregs expressing the 3A10 with CD25r and CD25f linkers had significantly higher pSTAT5 when compared to Tregs transduced with orthoIL-2Rβ alone and the 3A10 constructs with the two GS linkers showed milder increases. Moreover, Thy1.1^-^ non-transduced Tregs in the same culture of all 3A10 constructs showed minimal increase of pSTAT5 (Fig. 3c, d). Tregs expressing all tethered 1G12 constructs had increased pSTAT5 in Thy1.1^+^ and some showed increased pSTAT5 in Thy1.1^-^ Tregs (Fig. 3c, d). Lastly, wtIL-2-GS3 and GS4, but not the CD25- based linkers, showed increased pSTAT5 in Thy1.1^+^ and Thy1.1^-^ Tregs (Fig. 3c d). Thus, despite physically linking the IL-2 to its receptor, the tethered 1G12 and wtIL-2 could signal in neighboring non-engineered wtIL-2R-expressing cells depending on the linker used. Tregs engineered with 3A10-CD25r and 3A10-CD25f were significantly enriched between days 1 and 5 after IL-2 withdrawal (Fig. 3e), likely due to increased Bcl-2 expression (Fig. 3f, Fig. S7c) and more active proliferation (Fig. 3g, Fig. S7d). These *in vitro* experiments identified 3A10 tethered to orthoIL-2R with the CD25 linkers as promising candidates for selective and cell-autonomous support of Treg survival and proliferation.

### Transcriptomic program of Tregs expressing tethered orthoIL-2-IL-2R

The finding that the weak agonist, 3A10, induced stronger IL-2 signaling when tethered to the IL-2R was unexpected. We thus performed bulk RNAseq of 3A10-CD25r- and 1G12-CD25r- transduced Tregs (abbreviated to 3A10t and 1G12t hereafter) in the absence of IL-2 to explore potential explanations for their distinct behaviors. EV-transduced Tregs with or without IL-2 were included as controls. Principal component analysis (PCA) revealed EV Tregs with IL-2 were most distinct and separated from others along the principal component (PC) 1 axis. The remaining samples separated along the PC2 axis with 1G12t and EV no-IL-2 Tregs clustered closer together (Fig. 4a).

**Fig. 4.**
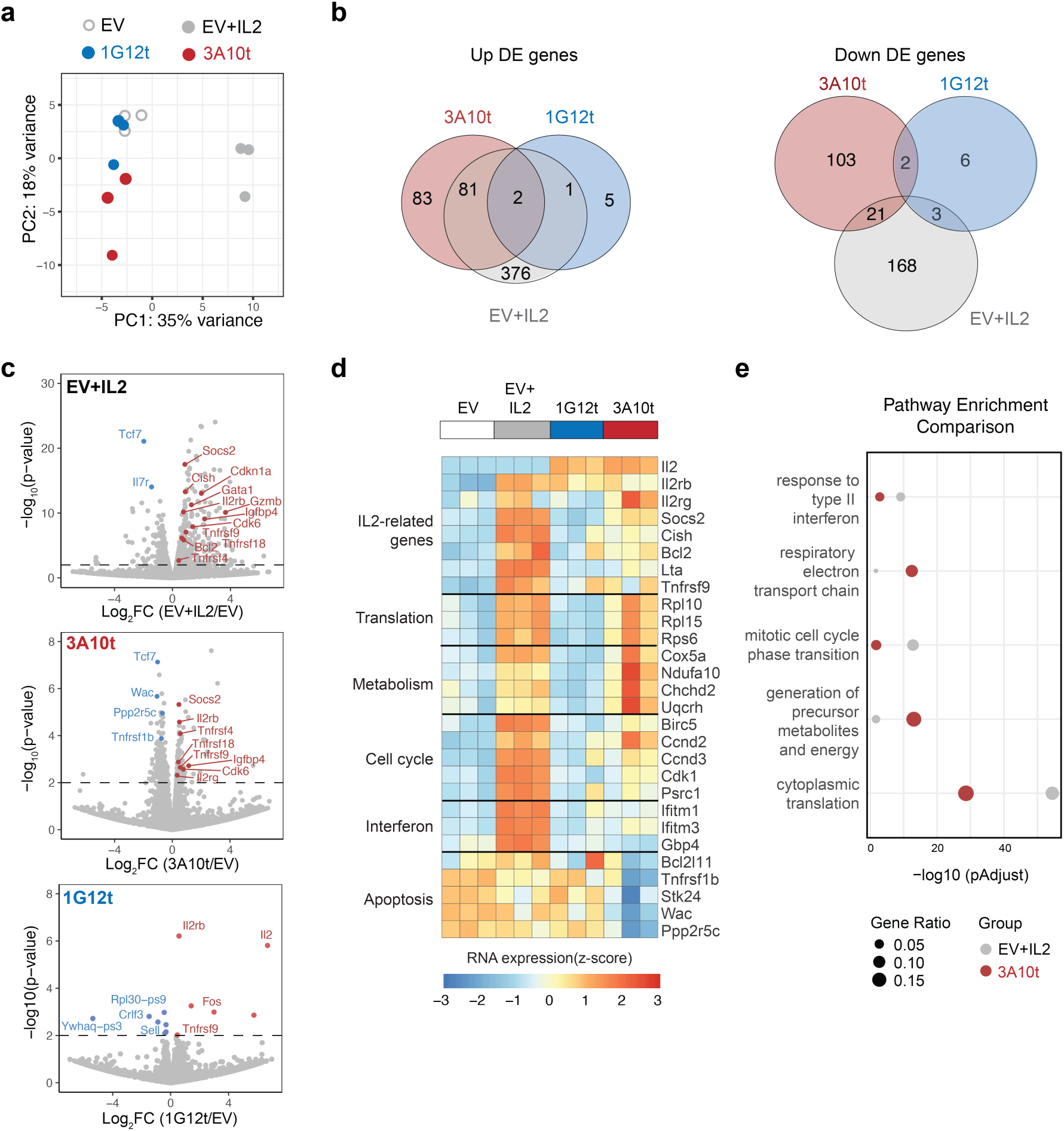
RNA-seq analysis reveals common and unique features of 3A10 Tregs. **a.** Principal-component analysis of RNA-seq data of Treg cells expressing empty vector (EV) with or without IL-2, and Treg cells expressing 1G12-CD25r or 3A10-CD25r. (**b.**) Venn diagrams showing overlap of upregulated (left) and downregulated (right) genes in EV+IL-2, 1G12-CD25r^+^ and 3A10-CD25r^+^ Treg samples when compare with EV without IL-2 samples. **c.** Volcano plots of differentially expressed genes (DEGs) in EV+IL-2 (top), 3A10- CD25r (middle), and 1G12-CD25r (bottom) Tregs when compared with EV. Horizontal dashed lines indicate P value of 0.01. Please refer to Data Table 1 online for a complete list of DE genes. **d.** Heatmap displaying expression of selected genes in EV, EV+IL-2, 1G12-CD25t+, and 3A10-CD25t+ Treg cells, grouped by pathways. **e.** Gene Ontology (GO) pathway comparison analysis showing the top 5 featured pathways among genes significantly upregulated in EV+IL-2 and 3A10-CD25t+ Treg cells. The plot depicts the adjusted p-values of DEGs in each pathway, with circle size indicating the gene ratio of DEGs within the pathway and color representing the different samples.

Pair-wised comparison with EV no-IL-2 revealed hundreds of differentially expressed genes (DEGs) in EV-IL-2 and 3A10t Tregs, but 1G12t had very few DEGs (Fig. 4b, c, Fig. S8a-c). Two of the upregulated genes in 1G12t Tregs were *IL2Rβ* and *IL2* (Fig. 4c, Data file S1), demonstrating successful transduction of these samples. Notably, both EV+IL-2 and 3A10t, but not 1G12t, has increased expression of *Socs2* and *Cish,* known negative feedback regulators of IL-2 signaling. These data suggested that the low signaling in 1G12t Tregs was likely not due to strong inhibition of IL-2 signaling.

We then compared the DEGs in EV+IL-2 and 3A10t groups to explore differential transcriptomic impact of soluble versus tethered IL-2 signaling in Tregs. Both groups showed increased IL-2 signaling indicated by upregulation of *Socs2, Cish, Bcl2, Lta, and Tnfrsf9* expression (Fig. 4d). Pathway analysis of the up-regulated DEGs in the two groups revealed shared cellular programs in cytoplasmic translation (*Rpl10, Rpl15, Rps6*). 3A10t showed stronger enrichment for metabolic genes (*Cox5a, Ndufa10, Chchd2, Uqcrh*), whereas EV+IL-2 had stronger activation of cell cycle (*Birc5, Ccnd2, Ccnd3, Cdk1, Psrc1*) and type II interferon response genes (*Ifitm1, Ifitm3, Gbp4*) (Fig. 4d, e, Fig. S8d). Selectively downregulated genes in 3A10t Tregs were enriched in apoptosis- and cell-cycle checkpoint-pathways (*Bcl2l11, Tnfrsf1b, Stk24, Wac, Ppp2r5c*) (Fig. 4e, Fig. S8e). Together, these transcriptomic analyses suggested that 3A10t promoted mitochondrial metabolism and suppressed apoptosis in Tregs, which might explain its efficacy in sustaining Tregs in the absence of IL-2.

### Tethered orthoIL-2 3A10 endowed Tregs autonomy from exogenous IL-2 and superior suppressive activity

To determine if autocrine signaling through 3A10t would alter the ability of 3A10t-expressing Tregs to sense and consume exogenous IL-2, we measured pSTAT5 signaling in response to titrated concentrations of IL-2 (Fig. 5a). Dose response of 3A10t-engineered Tregs was similar to that of EV-transduced and orthoIL-2R-transduced Tregs (Fig. 5b) and similar *in vitro* suppression when compared to Tregs expressing the orthoIL-2R alone (Fig. 5c).

**Fig. 5.**
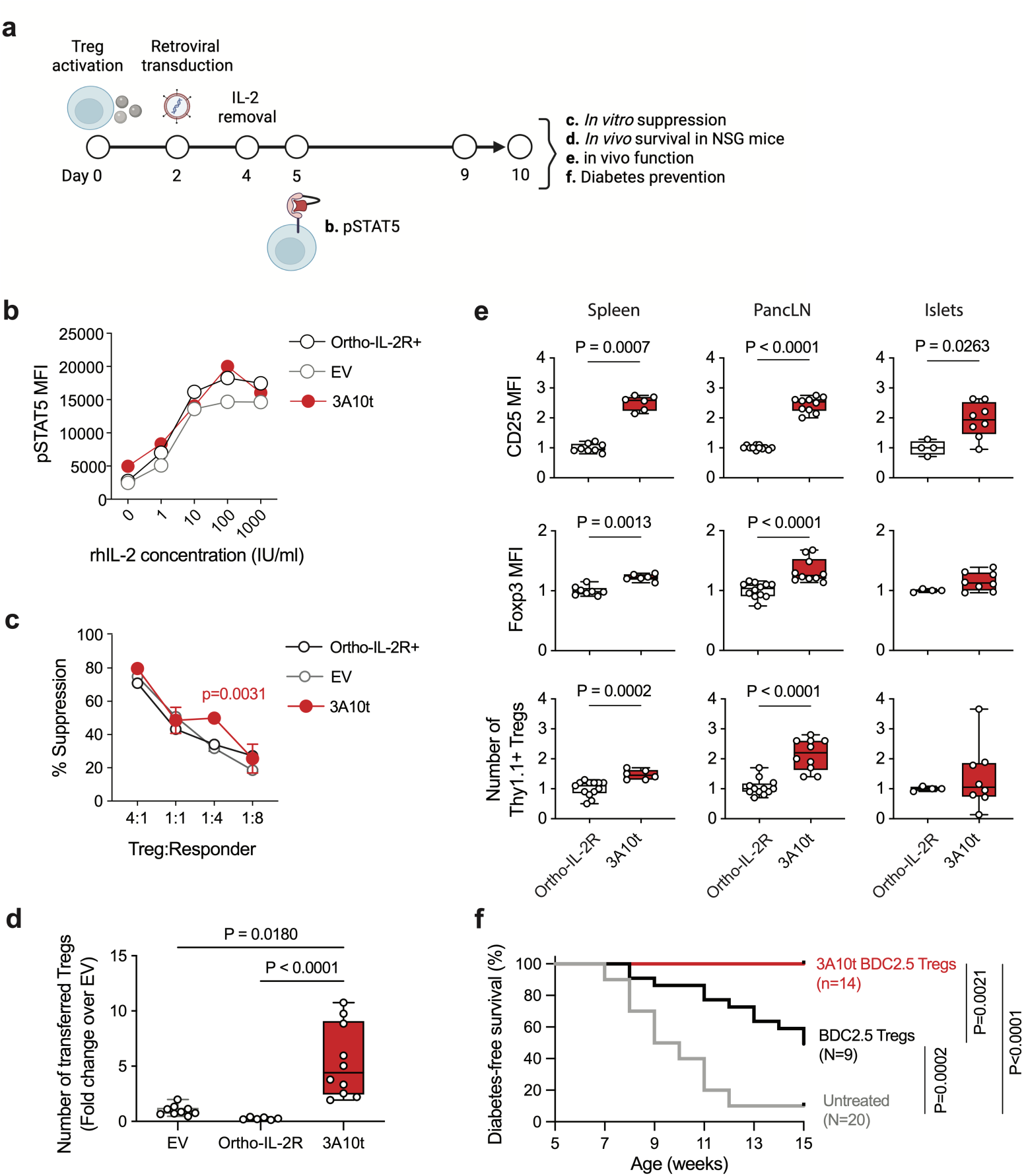
Function of Tregs expressing tethered 3A10-orthoIL-2R. **a.** Experimental workflow for assessing function of 3A10 expressing Treg *in vitro* and *in vivo.* **b.** pSTAT5 in response to titrated concentrations of exogenous IL-2 in Tregs transduced with empty vector (EV), *ortho*IL-2R, or 3A10t. Results shown are representative of 2 independent experiments. **c.** *In vitro* suppression. Ordinary two-way ANOVA followed by Tukey’s multiple comparison post-test was used to determine the statistical significance of the difference observed. P values on the graph are for comparisons between 3A10t^+^ and orthoIL-2R^+^ Tregs at each Treg:Responder ratio. Only P values <0.05 are listed. **d.** NSG mice were injected with Tregs transduced with EV, orthoIL-2R, or 3A10t. The numbers of transferred Tregs in the spleens of the recipient mice were analyzed 1 week after cell injection. Numbers shown were normalized to EV control. Data shown were a summary 3 independent experiments (means +/- SD, n=8-10 mice per group). Kruskal-Wallis test was used to determine the statistical significance among the groups. Only statistically significant P values are listed. **e.** NOD.CD28KO mice were injected with FACS-purified BDC2.5 Tregs transduced with orthoIL-2R or 3A10t. CD25 and Foxp3 expression in the transferred Treg and the numbers of transferred Tregs in the spleens, pancreatic LN (LN), and pancreatic islets were analyzed 1 week later. Data shown are a summary of 3 independent experiments (n=4-12 mice per group). Mann-Whitney test was used to determine the statistical significance between the two experimental groups. Only statistically significant P values are listed. **f.** NOD.CD28KO mice were treated with either 2,000 BDC2.5 + PBS or 3A10t Tregs. Mice with two readings of blood glucose >250 mg/dL were considered diabetic. (n=14-28 mice per group. 2 experimental replicates). Statistical significance was determined using Kaplan-Meier survival analysis. P values were calculated using Mantel-Cox test comparing all experimental groups.

A significantly higher number of 3A10t-transduced Tregs were recovered one week after transferring to NSG mice when compared to EV- or orthoIL-2R-transduced Tregs (Fig. 5d). In NOD.CD28KO mice, we observed increased CD25 and Foxp3 expression on 3A10t-engineered BDC2.5Tregs and increased cell number in the spleens and pancLN (Fig. 5e). BDC2.5 Tregs transduced with 3A10t had significantly higher CD25 expression than orthoIL-2R-transduced Tregs in the islets (Fig. 5e). A single infusion of 2,000 3A10t-engineered BDC2.5 Tregs completely prevented diabetes in NOD.CD28KO mice (Fig. 5f).

### Knocking tethered orthoIL-2 into Foxp3 locus safeguard against toxicity of off-Treg expression

Our results thus far showed that a tethered orthoIL-2-IL-2R system could sustain Tregs in the absence of exogenous IL-2. To determine if this system could also potentiate proinflammatory Tconvs, we expressed 3A10t in islet-specific CD4^+^ Tconv and assessed their diabetogenic potential in young prediabetic NOD.CD28KO mice. BDC2.5 Tconv cells expressing the 3A10t construct accelerated diabetes development when compared with empty vector-transduced BDC2.5 Tconvs (Fig. 6a).

**Fig. 6.**
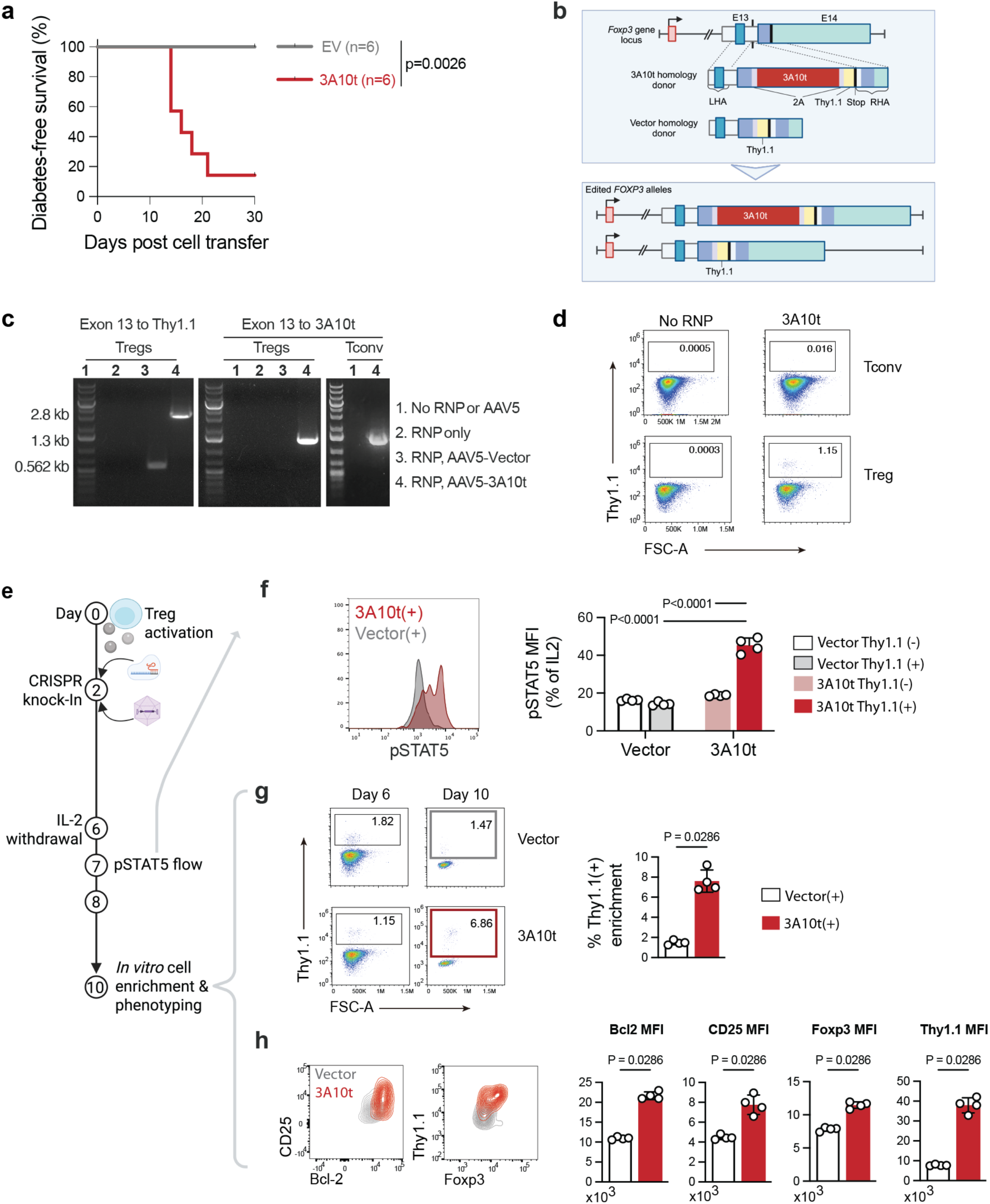
**Safe engineering of the tethered orthoIL-2-IL-2R system**. **a.** Diabetes development in NOD.CD28KO mice that received 20,000 CD4^+^ BDC2.5Tg TCR transgenic T cells transduced with empty vector (EV) or 3A10t. Results shown are a summary of 2 independent experiments. Statistical significance was determined using Kaplan-Meier survival analysis. P values were calculated using Mantel-Cox test. **b.** Schematic of 3A10t and vector knock-in strategy. (**c.**) PCR of genomic DNA showing vector and 3A10t insertion into the Foxp3 locus. **d.** Flow cytometric analysis of Thy1.1 expression in Tregs and Tconvs 4 days after editing. (**e.**) Experimental workflow for evaluating the function of 3A10t Foxp3 knock-in in NOD Tregs. **f.** pSTAT5 1 day after IL-2 withdrawal. **g.** Fold change in number of Thy1.1^+^ Tregs cultured without IL-2. **h.** Expression of Bcl2, CD25, Foxp3, and Thy1.1 by Thy1.1^+^ cells in vector knock-in and 3A10t knock-in Tregs cultured without IL-2. Statistical significance was determined using Mann Whitney test. P values for statistically significant differences between the vector and 3A10t groups are listed.

To ensure the selective expression of the orthoIL-2-IL-2R system in Tregs, we targeted the 3A10t construct to the Foxp3 locus by inserting it in the last intron of the *Foxp3* gene (Fig. 6b). PCR of genomic DNA from edited cells showed successful knock-in of the vector and the 3A10t constructs in both Tregs and Tconvs (Fig. 6c). Thy1.1 could be detected on the edited Tregs, not on Tconvs, demonstrating Treg-selective expression of the construct (Fig. 6d).

To determine if the expression of 3A10t driven from the Foxp3 locus could support Tregs in the absence of exogenous IL-2, we removed IL-2 from the Treg cultures and tested the cells using a battery of *in vitro* assays (Fig. 6e). Tregs with 3A10t knock-in exhibited higher pSTAT5 MFI (Fig. 6f) and showed selective survival advantage after IL-2 withdrawal (Fig. 6g). 3A10t knock-in Tregs had higher expression of Bcl2, CD25, and Foxp3 (Fig. 6h). Intriguingly, we observed significantly higher Thy1.1 expression on 3A10t knock-in Tregs, but not in vector knock-in Tregs, suggesting a positive feedback loop of IL-2 signaling enhancing Foxp3 and 3A10t-Thy1.1 expression (Fig. 6h). These results show that knocking the 3A10t construct into the Foxp3 locus could achieve Treg-specific expression and functional support of Treg autonomy with the added benefit of positive feedback loop reinforcing autocrine receptor expression and Treg lineage identity.

## Discussion

Although Tregs constitutively express the high-affinity trimeric IL-2R complex, delivering IL-2 selectively to Tregs remains an elusive goal due to the narrow therapeutic window between targeting Tregs and activating other IL-2-responsive cells, such as NK and activated T cells. The orthogonal autocrine IL-2-IL-2R system developed in this study provided an effective solution to address this critical barrier to Treg therapy. This system sustained Treg persistence regardless of exogenous IL-2, which can be limiting in diseased settings(Tang et al., 2008; Yshii et al., 2022) (Driver et al., 2012; Yamanouchi et al., 2007; Yang et al., 2022). The enhanced proliferation and survival of Tregs expressing autocrine orthoIL-2-IL-2R allowed the infused Tregs to establish persistent numerical dominance over inflammatory cells. This approach also led to increased expression of CD25 and Foxp3, molecules important for Treg function and lineage stability.

The stark functional differences among different tethered IL-2 constructs were unexpected and informative of the rules for effective cell engineering. The similar transcriptomic profile between Tregs expressing 1G12t and Tregs deprived of IL-2 suggested a near complete lack of signaling by the tethered 1G12. This suggested that the block of 1G12t signaling might be upstream of gene expression instead of induction of negative regulators of IL-2 signaling. It is possible that 1G12 engaged its orthogonal receptor shortly after its synthesis which led to ligand-induced receptor endocytosis and degradation of 1G12t(Cendrowski et al., 2016; Chen et al., 2017). This process may be less efficient for the low-affinity ligand 3A10t. Another interpretation is epigenetic silencing in cells with strong chronic IL-2 signaling(Moro et al., 2022). Lastly, persistent strong IL-2 signaling through 1G12t might have resulted in activation-induced cell death(Green et al., 2003). Future investigation into differential signaling and epigenomic impact by different tethered IL-2-IL-2R may shed light on the molecular mechanisms underlie an effective tethered IL-2 construct for Tregs.

Additionally, this study raises the question regarding the broader impact of autocrine IL-2 signaling in Tregs. Although it is generally accepted that Tregs do not produce IL-2, thus only rely on paracrine IL-2 signaling, this dogma has been challenged in a recent report suggesting a role of Treg-intrinsic IL-2 in Treg generation and maintenance(Chawla et al., 2020). A previous report in CD8^+^ T cells showed autocrine IL-2 induced weaker STAT5 signaling that limited effector differentiation in favor of a memory cell fate(Kahan et al., 2022). Pre-assembly of the IL-2R complex can occur in the ER and Golgi in autocrine cells(Volko et al., 2019), thus autocrine IL-2-IL-2R signaling could be initiated intracellularly whereas paracrine signaling could only initiate at the cell surface. This distinction may lead to differential engagement of signaling pathways that are responsible for distinct cellular behaviors. Constructs generated in this study will be useful for investigating this new aspect of Treg biology.

Taken together, this study demonstrates that it is possible to engineer an orthogonal autocrine IL-2-IL-2R system to improve the efficacy of therapeutic Tregs. This autocrine system has the advantage of being cell autonomous and not reliant on the exogenous provision of a ligand. In addition, this study reveals a safe and effective design as well as the importance of the model system for evaluating this cell engineering concept. These design and testing principles can help guide the development of an autocrine IL-2 system for human Tregs for clinical translation.

## Materials and methods

### Animals

Female and male NOD/ShiLtJ (Jackson laboratories), NOD.Thy1.1 & NOD.Thy1.2, NOD.Foxp3 cre/eGFP (Zhou et al., 2008), NOD.CD28KO (Salomon et al., 2000), NOD.BDC2.5 TCRTg, and NOD-scid IL-2Rgamma^null^ (NSG) mice were housed and bred in accordance with the UCSF (San Francisco, CA). Institutional Animal Care and Use Committee guidelines. Littermates were used as controls when possible, and all animals were age-matched in diabetes studies. NOD.Thy1.2 and NOD.Thy1.1 mice, 7-14 weeks, were used as donors of Treg and naive CD4^+^ T cells. NSG mice were used as recipients at ages 7-14 weeks for experiments. NOD.CD28KO mice were used as recipients in experiments starting at 5-6 weeks of age. Mice used in experiments were randomized based on age and sex so that these variables are equally distributed among experimental groups. All mouse experiments were performed according to a UCSF Institutional Animal Care and Use Program (IACUP)-approved protocol (IACUC protocol no. AN200668).

### Cell culture media

Cells were cultured in DMEM-high glucose no glutamine media supplemented with 10% heat-inactivated fetal calf serum, 100 IU/ml penicillin and streptomycin, 10mM of HEPES, GlutaMax, sodium pyruvate, non-essential amino acids, and 50 μM β-mercaptoethanol. Cells were resuspended in PBS supplemented with 2% heat inactivated fetal calf serum for isolation, enrichment, and sorting. Vendor and catalog information for key cell culture reagents can be found in Supplemental Table S1.

### Mouse Treg isolation and expansion

Single cell suspensions were prepared from lymph nodes (cervical, mediastinal, axillary, inguinal, mesenteric pancreatic) and spleen of the indicated NOD mice. CD4^+^ cell enrichment was carried out with EasySep mouse CD4^+^ T cell isolation kit. The enriched cells were stained with CD4, CD8, CD62L and CD25. Tregs were purified using florescence activated cell sorting (FACS) as CD4^+^CD62L^+^CD25^+^ or CD4^+^CD25^+^Foxp3.GFP^+^ and naive CD4^+^ T cells, as CD4^+^CD62L^+^CD25^-^ or CD4^+^CD25^-^ Foxp3^-^. Flow cytometric reactions and antibodies used are summarized in Supplemental Table S2. Cells were isolated using BD Aria2 SORP S854.

FACS-purified Tregs were stimulated with CD3/CD28 dynabeads at the ratio of 3:1 (beads:Treg) and cultured in cell culture media supplemented with 2,000 IU/ml rhIL-2. FACS-purified conventional T cells were stimulated at 1:1 bead to cell ratio and cultured in cell culture media supplemented with 200 IU/ml rhIL-2. The cells were counted and split every 2-3 days until utilized in an assay.

### Analytical flow cytometry

For STAT5 phosphorylation, transduced Treg cultures were washed on day 4 after activation to remove rhIL-2 in the cultures and rested for 18 hours in culture medium without IL-2. Varying concentrations of wildtype recombinant mouse IL-2 (wtIL-2) or orthoIL-2 (1G12 or 3A10) were then added to the cells. The cells were incubated for 30 minutes at 37°C. The cells were fixed with 4% PFA for 10 minutes and permeabilized with 100% ice-cold methanol for 30 minutes.

Cells were stained with anti-pSTAT5 AF647 and anti-Thy1.1 BV605. pSTAT5 mean fluorescence intensity (MFI) on Thy1.1^+^ cells were measured using flow cytometry. Cells were stained for extracellular surface markers for 30 minutes on ice in FACS buffer, fixed with a Foxp3 transcription factor staining buffer set for 30 minutes on ice, permeabilized and stained for intracellular markers for 30 minutes to 1 hour on ice. Information for all antibodies used are summarized in Supplemental Table S2. All samples were run on a BD LSRII.

### Retroviral constructs and T cell transduction

Retroviral constructs were cloned into an MSCV vector contain a mouse Thy1.1 reporter behind an internal ribosomal entry site (IRES). The orthoIL-2R and various tethered IL-2-IL-2E constructs were cloned upstream of the IRES. Retroviral Plat-E packaging cells were transfected using lipofectamine 2000 with the following plasmids: MSCV.IRES.Thy1.1 (empty vector), MSCV.orthoIL-2R.IRES.Thy1.1, or MSCV.orthoIL-2.Linker.orthoIL-2R.IRES.Thy1.1 and.

Virus was collected 48 hours later and used fresh or concentrated using Retro-X concentrator. Mouse Tregs were transduced on day 2 after activation with fresh virus and 20 μg/ml of polybrene. 200,000-300,000 Tregs were spinfected for 90 minutes at 25°C 600g. Following spinfection, virus was removed, and cells were resuspended in culture media supplemented with 2,000 IU/ml rhIL-2. Key molecular biology reagents used are summarized in Supplemental Table S3. Amino acid sequences of all constructs are listed in Supplemental Table S4.

### *In vitro* T cell proliferation assay

NOD Tregs were expanded in vitro for 10 days, washed with cell culture media to remove exogenous IL-2, and rested for 18 hours. Cells were stained with Cell Tracker Violet (CTV, Table S1) and restimulated with CD3/CD28 dynabeads (1:1). Cells were assessed 72-96 hours later for CTV dilution.

### *In vitro* suppression assay

Transduced Tregs from NOD.Thy1.2 mice were FACS purified based on Thy1.1 expression on day 10 after activation and rested for 6 hours. Naive CD4^+^CD62L^+^CD25^-^ cells from NOD.Thy1.1^+^ T cells were FACS purified and stained for CTV. Thy1.2^+^Thy1.1^+^ Tregs and naive Thy1.2^-^Thy1.1^+^ responder CD4 T cells were co-cultured at 4:1 1:1, 1:4, 1:8 and stimulated with plate-bound 0.5 μg/ml anti-CD3 and 1 μg/ml anti-CD28.

### *In vivo* experiments

Tregs were resuspended in 100 μl PBS and injects into NSG or NOD.CD28KO recipients retro-orbitally. orthoIL-2 and IL-2 were injected intraperitoneally in 100 μl volume. Diabetes development in NOD.CD28KO mice was assessed by measuring blood glucose. Mice with two readings of blood glucose higher than 250 mg/dL were considered diabetic.

### Tethered IL-2-IL-2R structure modeling

The predicted structure of tethered orthoIL-2-IL-2R was generated with AlphaFold protein structure databases. The predicted structure was then modeled with PyMol, a molecular visualization software (https://www.pymol.org/). Each protein is colored differently for recognition.

### RNA sequencing

Purified mouse Tregs were transduced with either 1G12t, 3A10t, or EV as a control on day 2 after activation. The cells were washed to remove exogenous IL-2 on day 4 (2 days after transduction) and cultured for 5 additional days. Half of the EV transduced culture was left in IL- 2-containing medium as a positive control. The transduced cells were FACS purified based on the expression of the Thy1.1 on day 9. Cells were lysed using TCL buffer (Qiagen), and RNA was extracted using VAHTS RNA Clean Beads (Vazyme), then eluted in 10µL of nuclease-free water. cDNA synthesis and amplification were performed using the Discover-sc WTA kit V2 (Vazyme) according to the instruction provided by the manufacture. The amplified cDNA products were purified using VAHTS DNA Clean Beads (Vazyme) and quantified using Qubit. Illumina sequencing libraries were made using 50 ng of the amplified cDNA using the TruePrep DNA Library Prep Kit V2 for Illumina (Vazyme) and the TruePrep Index Kit V2 for Illumina (Vazyme). Size selection of the PCR product was performed using 0.55x VAHTS DNA Clean Beads. Finally, 20 ng of each sample were pooled for subsequent second-generation sequencing on the Illumina NovaSeq X, with a read length of 50 bp for paired end. Key molecular biology reagents used are summarized in Supplemental Table S3.

Raw fastq reads were trimmed by Cutadapt (version 1.18) to trim adapter and low-quality sequence. Reads were aligned to the mouse genome (mm10) using STAR (version 2.5.3). The number of reads within each gene in each sample were counted using RSEM (version 1.3.0) with gene annotation file from GENCODE (GRCm38.m23). Differential expression was estimated by using DESeq2 package (version 1.26). Differentially expressed genes in EV+IL-2, 3A10-CD25t and 1G12-CD25t Treg in comparison to EV without IL-2 Tregs were defined as log2 Fold change > 0.25 and P value < 0.01. Pathway analysis was performed by using the cluster Profiler R package (v3.14.3) with default parameters (P value < 0.05) and the functional annotations terms in Gene Ontology (GO) (Yu et al., 2012).

### AAV production

The fragments LHA-oExon14-P2A-Thy1.1-Stop-RHA and LHA-oExon14-T2A-3A10t-P2A- Thy1.1-Stop-RHA were cloned into AAV2-ITR transfer plasmids. These transfer plasmids, along with pAAV2/5 Rep-Cap plasmids and adenovirus helper plasmids, were transfected into HEK293T cells using polyethylenimine to package the donor template into AAV5 capsids. On day 3 after transfection, cells were scraped and collected in AAV lysis buffer (50 mM Tris, 150 mM NaCl). The cells were lysed using 3 rounds of freeze and thawing, followed by a 1-hour incubation at 37°C with 25 units/mL Benzonase (Millipore Sigma #70-664-3). The AAV particles were purified using iodixanol gradient ultracentrifugation (OptiPrep, StemCell Technologies). The iodixanol layer between 40% and 54% was extracted using an 18-gauge needle. The AAV was washed and concentrated using storage buffer (1X PBS with 0.001% Tween-20).

### CRISPR knock in

Lyophilized gRNA (AGCCUGGGGCUAGACAUGUG) (IDT) was resuspended in nuclease-free TE buffer at a concentration of 100 μM, aliquoted, and stored at −80°C. Recombinant Cas9- NLS (40 μM) was purchased (QB3 Macrolab). For each reaction, a mixture of 1:1 ratio of gRNA and Cas9 (60 pmol: 60 pmol) was incubated at room temperature for 15 minutes to form ribonucleotide-protein complexes (RNP). Dynabeads were removed from activated Treg or Tconv cells on day 2 after activation using magnetic separation. For each electroporation reaction, 200,000 cells were resuspended in 20 μL Lonza electroporation buffer P3 and mixed with 3 μL of RNP, and electroporated in a Lonza 4D 96-well electroporation system using pulse code DN100 (Treg) or EH115 (Tconv). After electroporation, 80 μL of pre-warmed culture medium was immediately added to the electroporated cells and the cells were rested for 10 minutes at 37°C. The rested cells were seeded in wells of 96-well round-bottom plates in 200 μL culture medium and 20 μL AAV virus. Cells were subsequently cultured and maintained at a density of 1 million cells/mL.

### Statistics

Statistical analyses were performed with the GraphPad Prism 9 software. In diabetes protection experiments with prior knowledge of the incidence of untreated mice at 80-90%, power calculations were performed. In general, minimally 8 mice per experimental group is needed to detect a significant reduction of diabetes incidence by 50%. When no power calculation was possible due to a lack of prior data, 3 to 5 biological replicates were included per experimental condition and all results shown were repeated at least once in an independent experiment with similar results. Mann-Whitney test was used to determine the statistical significance between the two experimental groups. Ordinary one-way ANOVA Kruskal-Wallis test followed by Dunn’s multiple comparison test was used to determine statistical significance among 3 or more experimental groups with one independent variable. Ordinary two-way ANOVA followed by Tukey multiple comparison test was used to determine statistical significance among 3 or more experimental groups with two independent variables. Statistical significance of diabetes-free survival was determined using Kaplan-Meier survival analysis and using Mantel-Cox test to calculate p values comparing each condition to BDC2.5 Treg-treated group.

### Online supplemental material

Fig. S1 is a new analysis of previously published clinical data supporting the rationale of the current study. Figs. S2 to S4 contain illustrations and original data in support of Fig. 1. Figs. S5 and S6 contain illustrations and original data in support of Fig. 2. Fig. S7 contains modeling and original data related to Fig. 3. Fig. S8 contains additional data related to Fig. 4. Tables S1 to S3 contain detailed listing of key reagents used in the study. Table S4 contains amino acid sequences of constructs developed in this study. Differential gene expression data from RNAseq analysis is provided in Source Data file S1.

## Data availability

Data generated in this study are provided within the article itself and its supplementary materials.

## Acknowledgements

We thank M.S. Anderson, A. Marson, J. Eyquem, M.H. Spitzer, F.V. Gool, J.A. Smith, A. Young, L. Vo, P. Ho, W. Nyberg, G. Yuan, D. Simeonov, J.T. Cortez, I. Tenvoreen, D. Marquez, O. Aguilar, and the UCSF Parnassus Flow Cytometry CoLab for their support and advice. Figure panels 1a, 3a, 5a, 6b, 6e, S2a, S2b, S3a, S4a, S5b, and S6a were created in Biorender (BioRender.com/q07c632).

## Funding

NIH UC4 DK116264 (J.A.B., Q.T.), Sean Parker Autoimmune Laboratory fund (J.A.B.), Breakthrough T1D COE-2019-860-S-B (Q.T.), Klein-Kraft 2022-2023 Diabetes Research Fellowships (ACI), Graduate Division Travel Award (ACI), Diabetes, Endocrinology, and Metabolism NIH training grant T32 (ACI), IMSD UCSF Graduate Division Summer Research fellowship (ACI), Howard Hughes Medical Institute grant (KCG), Ludwig Institute grant (KCG), NIH RO1-AI51321 (KCG).

Author contributions: Conceptualization and Design of Study: A.C.I., H.L., J.A.B., Q.T., Methodology: A.C.I., H.L., J.A.B., Q.T., Investigation: A.C.I., H.L., V.Q.N., W.R., Y. C., Analysis: A.C.I., H.L., S.S.J., K.J., L.S., Q.T., Orthogonal IL-2 reagents and advice on experimental design: K.C.G., K.J., L.S., Writing-original draft: A.C.I., H.L., Q.T., Writing-review & editing: A.C.I., H.L., J.A.B., K.C.G. Q.T.

Disclosures: Q.T. is a cofounder and shareholder of Sonoma Biotherapeutics, holds shares of stock in eGenesis, and is a scientific advisor for Sonoma Biotherapeutics, Qihan Bio, Minutia, Moderna, and Waypoint Bio. Q.T. is a coinventor of the following relevant issued patents and pending patents: US9801911B2, US7722862B2, PCT/US2019/022546, PCT/US2019/023883, PCT/US2020/030869, PCT/US2021/072139, PCT/US2021/049682, PCT/US2022/074720, and US Patent Application No. 63/571,413. J.A.B. is CEO and president and has a financial interest in Sonoma Biotherapeutics and is on the board of directors and has financial interests in Gilead. J.A.B. is a coinventor of the following relevant issued patents and pending patents: PCT/US20230280341A1, PCT/US9801911B2, aU2016211161C1, PCT/US10138298B2, PCT/US7722862B2, PCT/US2019/022546, PCT/US2019/023883, PCT/US2021/072139, PCT/US2021/049682, and US Patent Application No. 63/571,413. K.C.G. is the founder of Synthekine Therapeutics, which has licensed the ortho-2 technology. K.C.G. is an inventor on a patent application describing the ortho-2 system (biologically relevant orthogonal cytokine/receptor pairs, US patent no. 10,869,887B2). K.C.G. and L.S. are shareholders of Synthekine, a biotechnology company that has licensed the ortho–IL-2 technology. All other authors have no conflict of interest to report.

