## Supplemental figures 1 to 8 for "Receptor-tethered orthogonal IL-2 enhances regulatory T cell therapy"

**Fig. S1. Pharmacokinetics of infused Tregs in the peripheral blood of type 1 diabetes patients.**

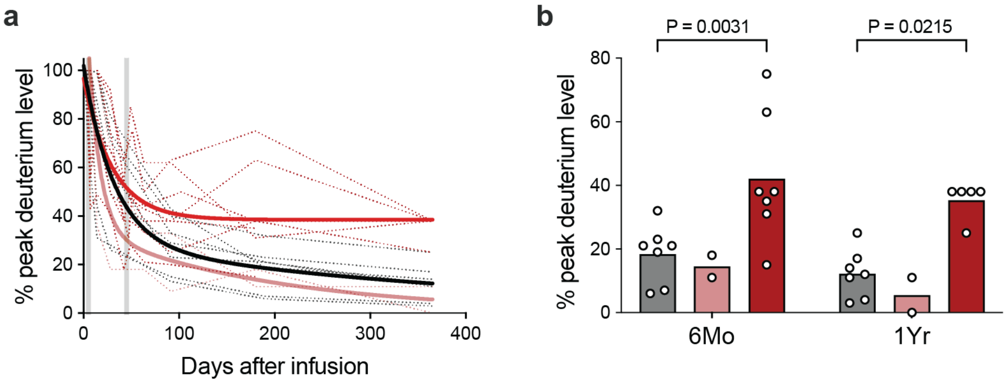


FACS-purified autologous Tregs were expanded ex vivo in medium containing deuterated glucose, which resulted in the enrichment of deuterium in the genome of the expanded Treg products. Percentage of deuterium enrichment in the peripheral blood Tregs can then be used to assess the pharmacokinetics of the Tregs post infusion. **a.** Deuterium signals in peripheral blood over time. The black lines are results from patients who received Tregs alone. The dark red lines are results from patients who received Tregs followed by two rounds of daily IL-2 infusion given between days 3-7 and 42-46. The light pink lines are results from patients who received Tregs followed by one round of daily IL-2 infusion given between days 3-7. The thin dotted lines are data for individual patients and the thick solid lines are the mean of the group. All data are normalized to peak deuterium enrichment detected between day 1 and 14 after cell infusion. **b.** Normalized deuterium enrichment at 6 month and 1 year after infusion in the 3 groups of patients were compared. Results shown are a re-analysis previously published data (DOI: 10.1126/scitranslmed.aad4134 and 10.1172/jci.insight.147474).

**Fig. S2. Experiments for evaluating orthoIL-2 3A10 on orthoIL-2R engineered Tregs** (related to main Fig. 2).

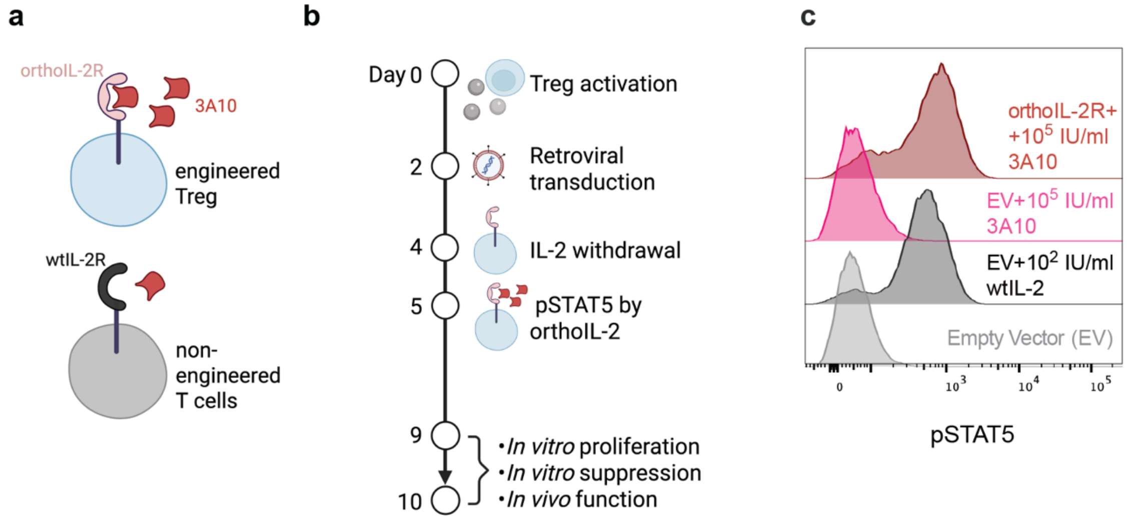


**a.** Schematic of the orthogonal IL-2-IL-2R system. **b.** Experimental workflow for engineering and testing the 3A10 and orthoIL-2R system. **c.** Representative flow cytometric histograms of phosphorylated STAT5 (pSTAT5) induction after 30-minute stimulation with 100IU/ml IL-2 or 100,000IU/ml orthoIL-2 3A10. Panels a and b were created in BioRender.

**Fig. S3. OrthoIL-2 3A10 mildly activated orthoIL-2R+ Tregs in NSG mice** (related to main Fig. 2).

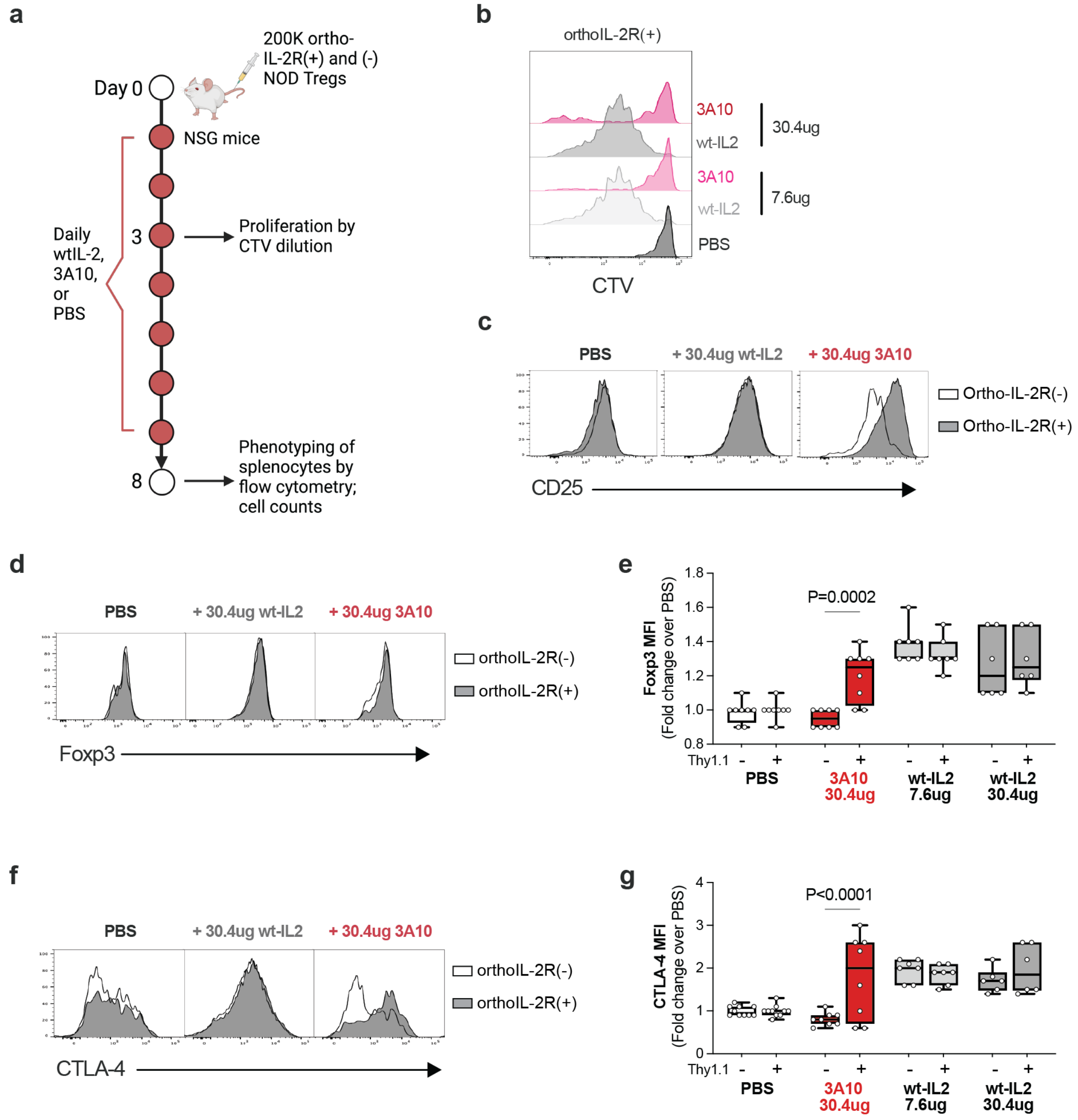


**a.** Experimental workflow for assessing 3A10 on orthoIL-2+ Tregs in NSG mice. **b.** Proliferation of orthoIL-2R+ Treg in NSG recipients. **c-g.** Splenocytes were collected on day 8 after Treg transfer and daily injection of the indicated reagents. Representative results of flow cytometric analysis of CD25 (**c**), Foxp3 (**d**), and CTLA4 (**f**) expression on transferred orthoIL-2R(+) and (-) Tregs are shown. Summary of flow cytometric analysis of Foxp3 (**e**) and CTLA-4 (**g**) expression are shown. Ordinary two-way ANOVA followed by the Sidak multiple comparison test was used to determine the statistical significance of the difference between orthoIL-2R (+) and (-) Tregs. P values for the data with significant differences are indicated on the graphs. Panels a was created in BioRender.

**Fig. S4. Weak activity of orthoIL-2 3A10 on orthoIL-2R+ Tregs in NOD.CD28KO mice** (related to main Fig. 2).

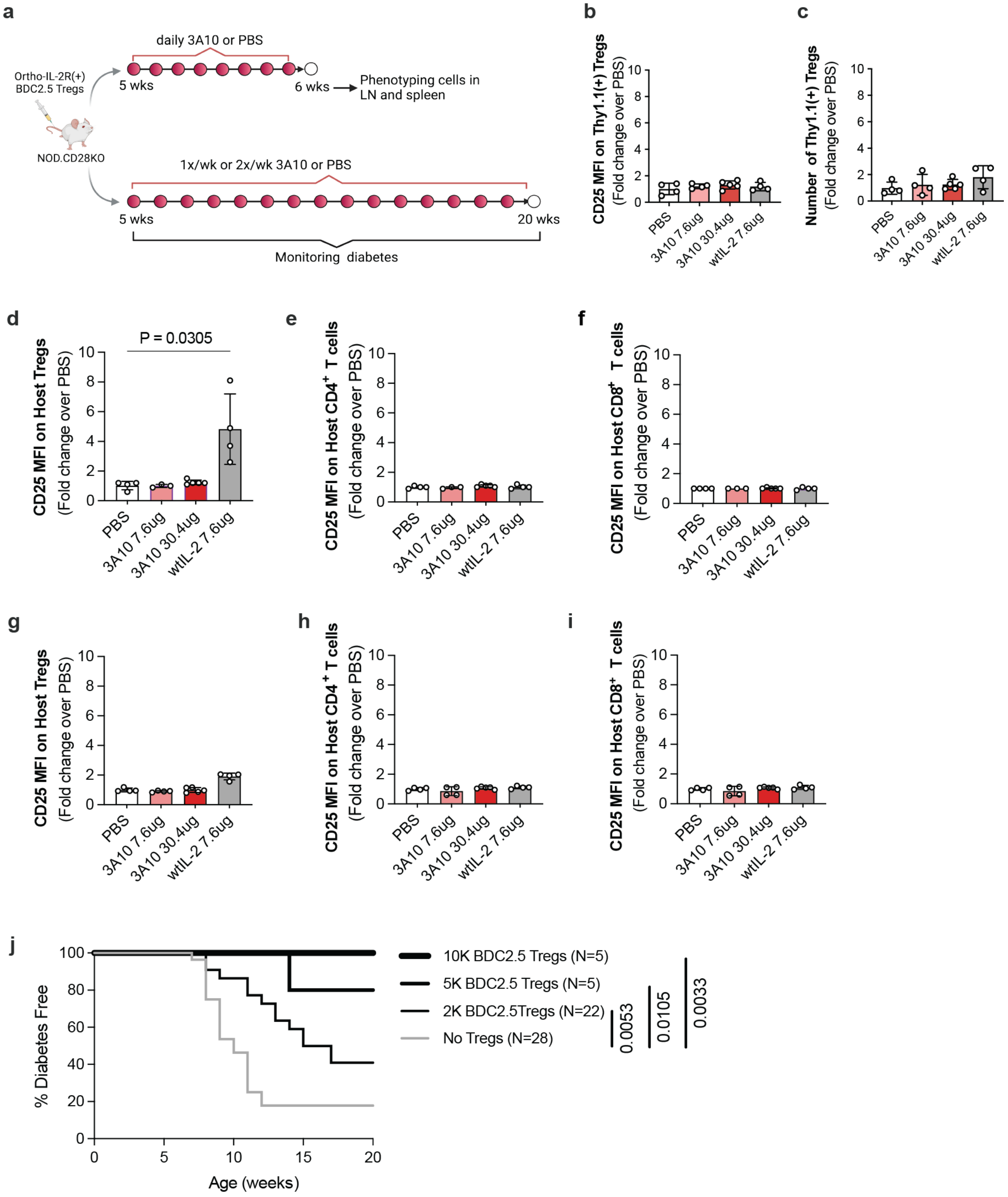


**a.** Experimental workflow for assessing 3A10 on orthoIL-2+ Tregs in NOD.CD28KO mice. **b, c.** effect of weeklong daily treatment with MSA-3A10 or MSA-wtIL-2 on orthoIL-2R+ Treg expression of CD25 (**b**) and total number (**c**). **d-i.** effect of weeklong daily treatment with MSA-

3A10 or MSA-wtIL-2 on recipient Tregs (**d, g**), CD4+ T cells (**e, h**), and CD8+ T cells in the spleen (**d-f**) and pancreatic LN (**g-i**). Results shown are summaries of 3-5 independent experiments. One-way ANOVA followed by the Kruskal-Wallis test was used to determine the statistical significance of the difference with PBS controls. **j.** A titrated doses of BDC2.5 Tregs were injected into 5-week-old NOD.CD28KO recipients to determine a sub-optimal dose for diabetes prevention. Mantel-Cox Log-rank test was used to determine the statistical significance of the differences in diabetes-free survival in comparison to the untreated controls. P value is only listed for significant differences. Panels a was created in BioRender.

**Fig. S5. Experiments for evaluating orthoIL-2 1G12 on orthoIL-2R engineered Tregs** (related to main Fig. 3).

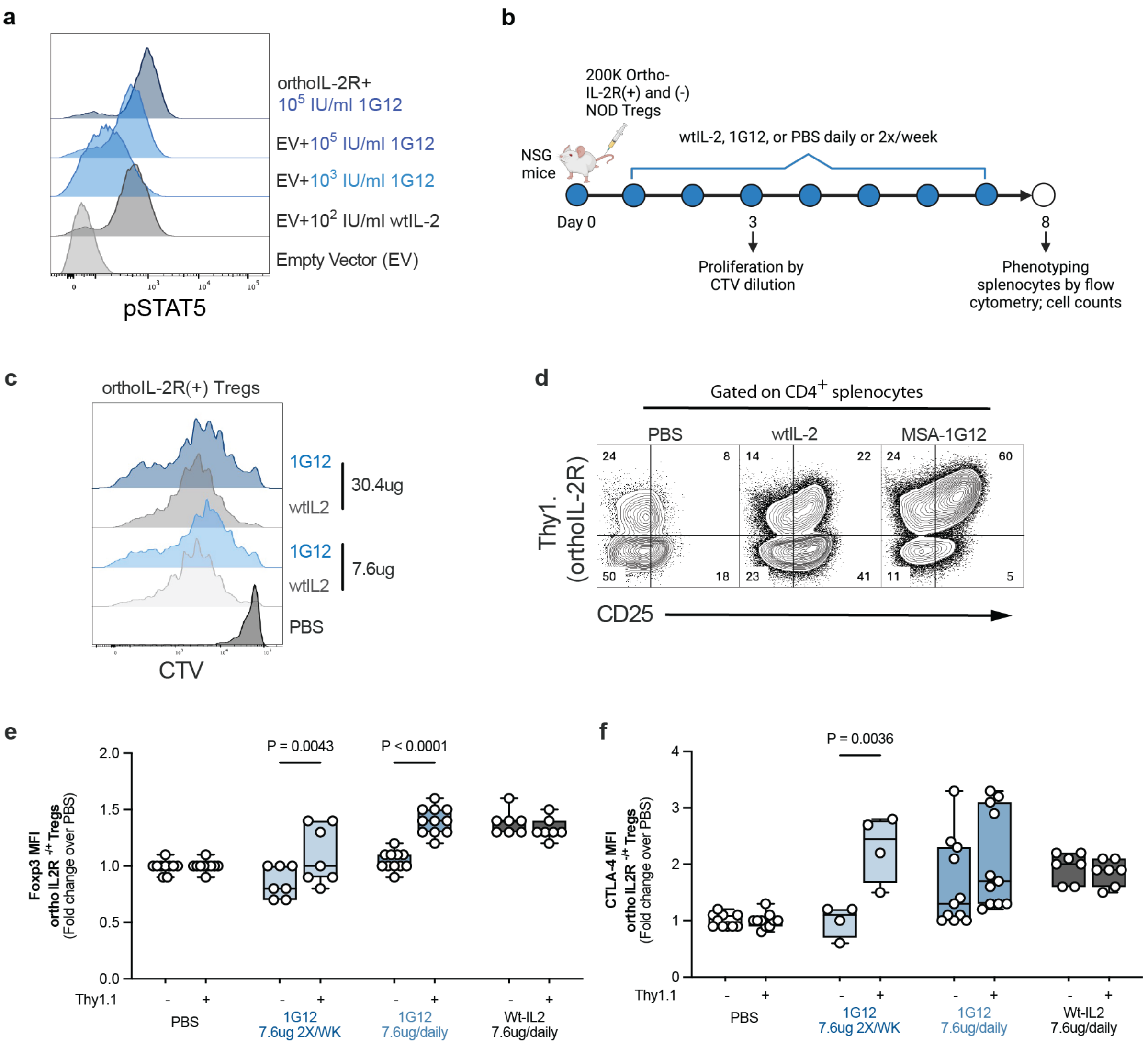


**a.** Representative flow cytometric profile of phosphorylated STAT5 induction after 30-minute stimulation with 100IU/ml wtIL-2 or 100,000IU/ml 1G12. **b.** Experimental workflow for assessing 1G12 in NSG mice. **c.** Representative flow cytometric plots showing in vivo proliferation of orthoIL-2R-transduced Tregs in NSG mice treated daily with 1G12 or wtIL-2. **d.** Representative flow cytometric plots showing CD25 versus Thy1.1 expression on transferred cells in the spleen. **e, f.** Summary of results of Foxp3 and CTLA4 expression on transferred orthoIL-2R (+) Th1.1 (+) and (-) Tregs in NSG mice receiving MSA-1G12 or MSA-wtIL-2 using doses indicated. Ordinary two-way ANOVE followed by the Sidak multiple comparison test was used to determine the statistical significance of the difference between orthoIL-2R (+) and (-) Tregs. Only significant P values are listed. Panel b was created in BioRender (BioRender.com/q07c632).

**Fig. S6. Activity of orthoIL-2 1G12 on orthoIL-2R+ Tregs in NOD.CD28KO mice** (related to main Fig. 3).

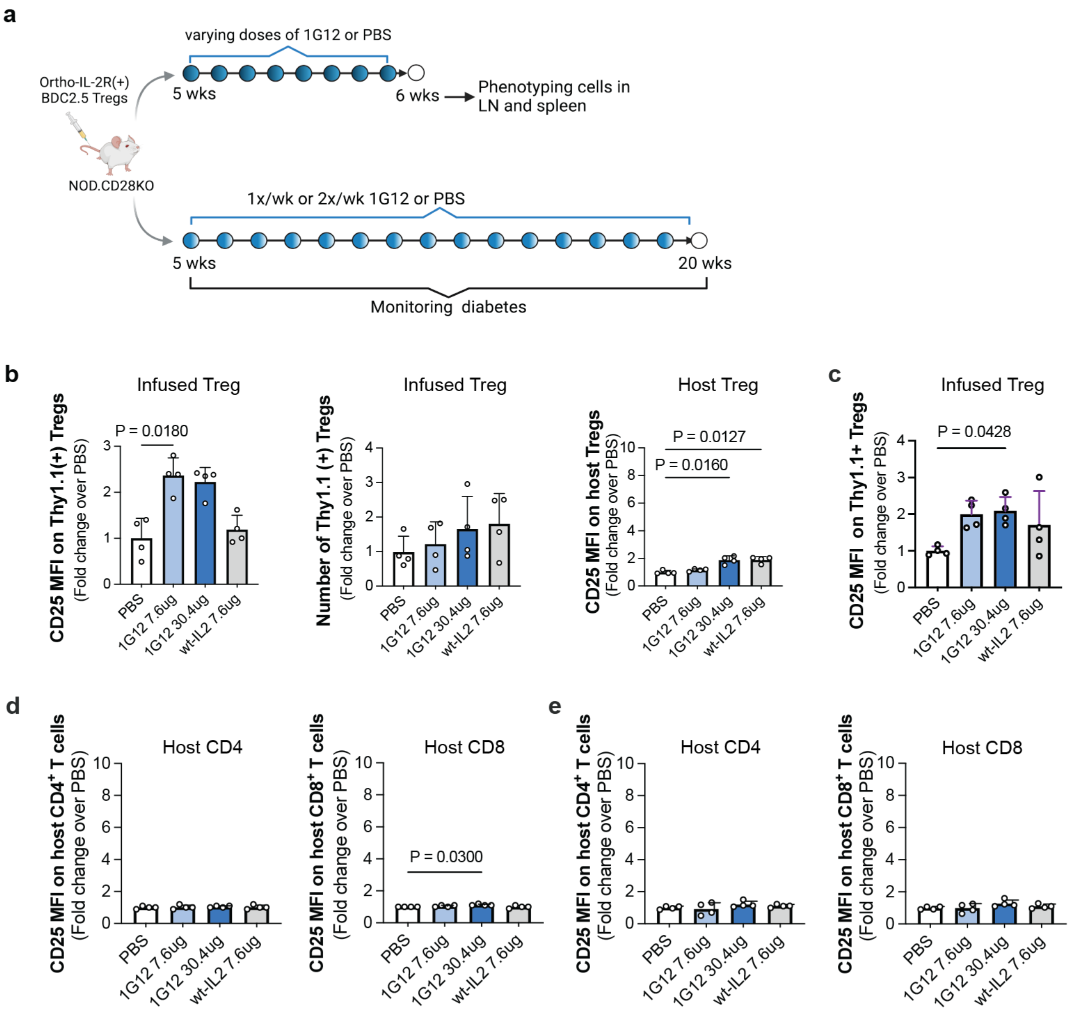


**a.** Experimental workflow using NOD.CD28KO mice. **b.** Infused orthoIL-2+ Tregs and host Tregs in the pancreatic LN after weeklong daily treatment with MSA-1G12 or MSA-wtIL-2 at the indicated dose. **c.** Analysis of CD25 expression on orthoIL-2R+ Tregs in the pancreatic islets of the recipient mice. **d, e.** Expression of CD25 on host CD4+ and CD8+ conventional T cells in the spleen (d) and pancreatic LNs (e). One-way ANOVA followed by the Kruskal-Wallis test was used to determine the statistical significance of the difference with PBS controls. Only significant P values are listed. Panel a was created in BioRender.

**Fig. S7. Structure prediction and function assessment of orthogonal IL-2-IL-2R constructs** (related to main Fig. 4).

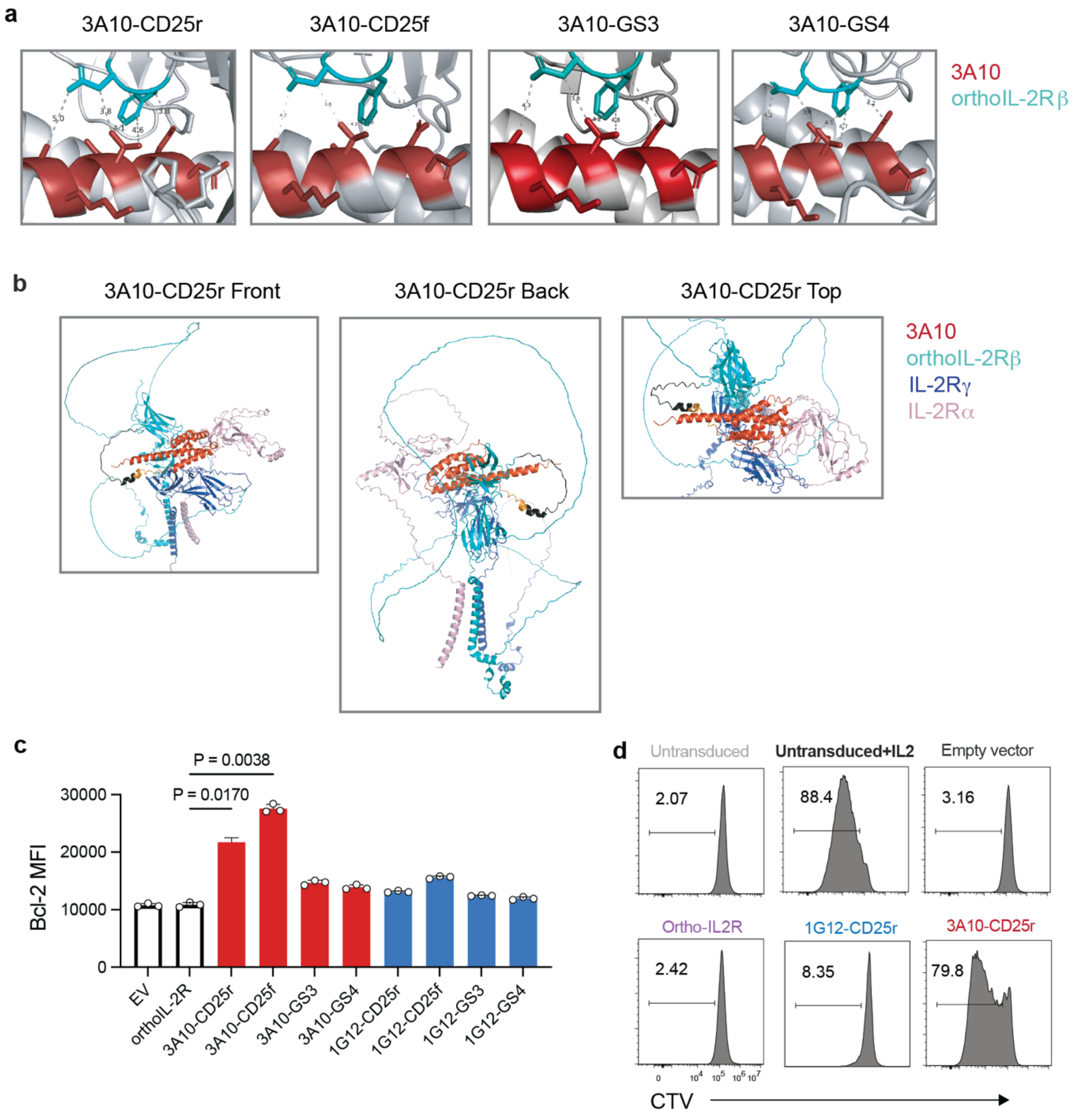


**a.** Alpha-fold models of the distance of 3A10's side chains from the orthoIL-2Rβ binding pocket when tethered to all four linkers. **b.** Alpha-fold models of the 3A10 and the trimeric IL-2R complex. **c.** Expression of Bcl-2 on day 5 3A10-CD25r- and 3A10-CD25f-expressing Tregs in the absence of exogenous IL-2. **d.** CTV dilution of 1G12-CD25r and 3A10-CD25r-expressing Tregs.

**Fig. S8. Distinct transcriptomic program in Tregs expressing various tethered orthoIL-2 constructs (**related to main Fig. 5).

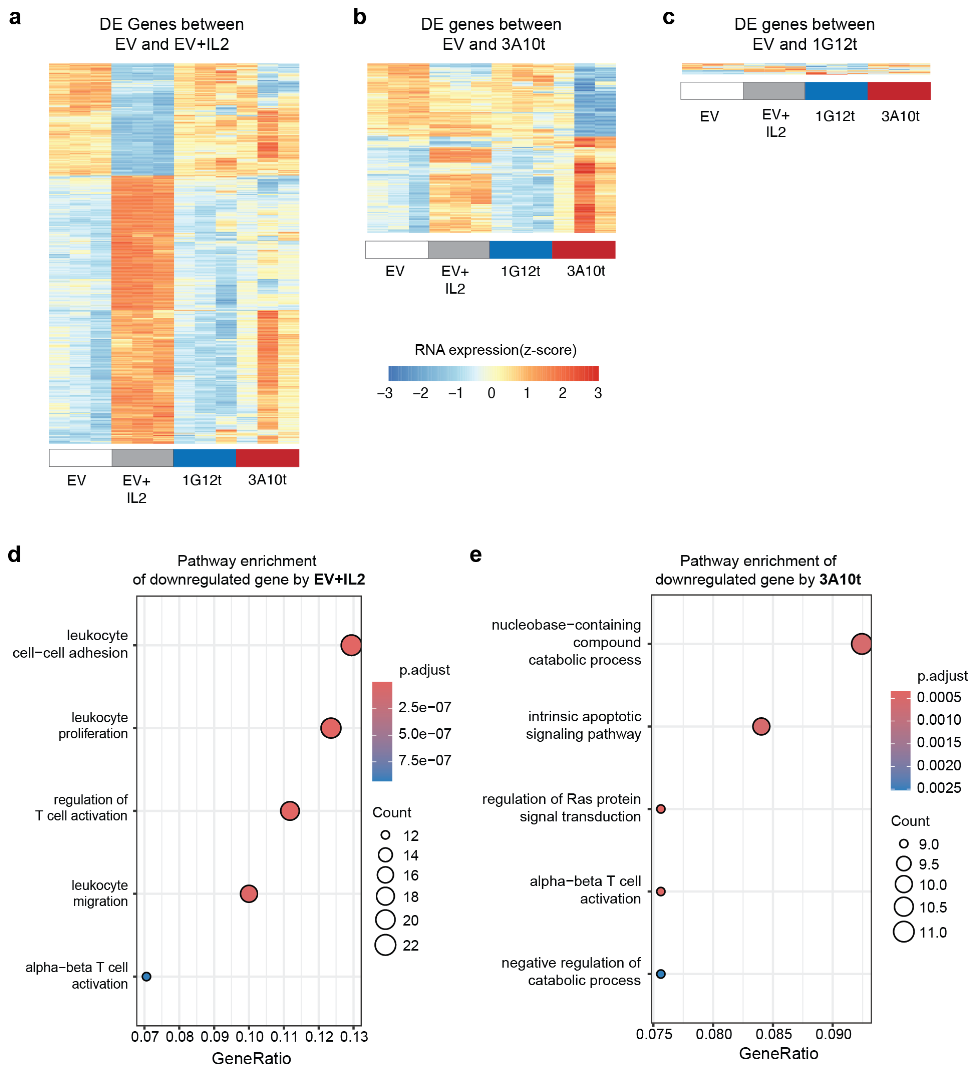


**a.** Heatmap of the expression of DE genes identified between empty vector (EV)+IL-2 and EV by all samples are shown. **b.** Heatmap of the expression of DE genes identified between 3A10t and EV by all samples are shown. **c.** Heatmap of the expression of DE genes identified between 1G12 and EV by all samples are shown. **d.** Gene Ontology (GO) pathway analysis showcasing the top 5 featured pathways among genes significantly downregulated in EV+IL-2 Treg cells when compared to EV Tregs. **e.** Gene Ontology (GO) pathway comparison analysis showcasing the top 5 featured pathways among genes significantly downregulated in 3A10t samples when compared to EV samples. In panels d and e, the plots depict the gene ratio of DE genes in each pathway, with the circle size indicating the gene count of DE genes in each pathway and the color representing the adjusted p-values.
